# Natural enemies have inconsistent impacts on the coexistence of competing species

**DOI:** 10.1101/2020.08.27.270389

**Authors:** J. Christopher D. Terry, J. Chen, O. T. Lewis

## Abstract

1. The role of natural enemies in promoting coexistence of competing species has generated substantial debate. Modern coexistence theory provides a detailed framework to investigate this topic, but there have been remarkably few empirical applications to the impact of natural enemies.
2. We tested experimentally the capacity for a generalist enemy to promote coexistence of competing insect species, and the extent to which any impact can be predicted by trade-offs between reproductive rate and susceptibility to natural enemies.
3. We used experimental mesocosms to conduct a fully-factorial pairwise competition experiment for six rainforest *Drosophila* species, with and without a generalist pupal parasitoid. We then parameterised models of competition and examined the coexistence of each pair of *Drosophila* species within the framework of modern coexistence theory.
4. We found idiosyncratic impacts of parasitism on pairwise coexistence, mediated through changes in fitness differences, not niche differences. There was no evidence of an overall reproductive rate – susceptibility trade-off. Pairwise reproductive rate – susceptibility relationships were not useful shortcuts for predicting the impact of parasitism on coexistence.
5. Our results exemplify the value of modern coexistence theory in multi-trophic contexts and the importance of contextualising the impact of natural enemies. In the set of species investigated, competition was affected by the higher trophic level, but the overall impact on coexistence cannot be easily predicted just from knowledge of relative susceptibility. Methodologically, our Bayesian approach highlights issues with the separability of model parameters within modern coexistence theory and shows how using the full posterior parameter distribution improves inferences. This method should be widely applicable for understanding species coexistence in a range of systems.

## Introduction

Species compete for limited resources and (in almost all cases) are consumed by other species. Understanding how the impacts of competitors and natural enemies interact to influence species coexistence is a long-standing challenge. Natural enemies are regularly cited as a driver of coexistence, for example by suppressing species that would otherwise be competitively dominant (Paine, 1966), but their impact can be highly variable (Chase et al., 2002).

Analyses investigating the effect of natural enemies often focus on ‘reduction in competition’ (e.g. (Gurevitch et al., 2000), as measured by negative effects between focal species; however consumers can affect competition and coexistence in convoluted ways (Chase et al., 2002; Pringle et al., 2019). Modern coexistence theory (MCT) focuses on the property of mutual invasibility (Grainger, Levine, et al., 2019), and allows the precise framing of questions regarding pairwise species coexistence as a balance between niche differences promoting stabilisation and fitness differences fostering exclusion of one species by another (Chesson, 2000; Letten et al., 2017). Through the lens of modern coexistence theory, interactions mediated through competition for resources and through natural enemies are conceptually equivalent (Chesson, 2018; Chesson & Kuang, 2008) and a body of theory has developed to integrate consumer effects within the coexistence framework (Chesson & Kuang, 2008; McPeek, 2019). This theoretical work has emphasised that impacts of any process on coexistence can only be determined with information on both intrinsic growth rates and interaction terms. Alongside the theory, empirical studies using the modern coexistence framework to investigate the impact of consumers on coexistence is now emerging (Bartomeus & Godoy, 2018).

Natural enemies can affect both fitness differences and niche differences between species. A frequently-invoked mechanism is that they reduce fitness differences, mediated by a trade-off between population growth rate and susceptibility to attack (Kraaijeveld & Godfray, 1997; Paine, 1966). However, such a trade-off is in itself insufficient to enable long-term coexistence, since some kind of stabilising niche-difference is required to mitigate any fitness difference (Chesson, 2000). Even with zero fitness difference, stochastic drift would preclude coexistence in the absence of stabilising mechanisms (Adler et al., 2007). Natural enemies can also potentially influence niche differences through modifying the overall competitive pressure exerted between competitors. Enemies may cause competitors to change their behaviour, or natural enemies may change their foraging behaviour as the relative abundances of the competing species change (Abrams, 2010).

Most experimental work within a formal MCT framework has been conducted on plant or microbial systems (Bartomeus & Godoy, 2018; Grainger, Letten, et al., 2019) including investigation of the effects of seed predation (Nottebrock et al., 2017; Petry et al., 2018), and seed pathogens (Mordecai, 2013). Experimental studies of insect communities have considerable potential to further our understanding of the role of natural enemies in species coexistence. Since parasitoids kill their insect hosts as they develop in or on them, each attack is directly interpretable in terms of host fitness (Hassell, 2000). Although ecologists have been competing insect species (especially *Drosophila*) against each other for decades (Davis et al., 1998; Gilpin et al., 1986; Pearl, 1932; Worthen, 1989), very few studies have explicitly tested for mutual-invasibility in insect systems (Siepielski et al., 2018), rather than particular conditions necessary for coexistence (Godwin et al., 2020; Spaak & de Laender, 2020).

Here we present an analysis parameterising the impact of consumers on pairwise species coexistence, focusing on a community of six rainforest *Drosophila* species. Our experiment was designed specifically to capture the key parameters of our focal species and their competitive interactions. We examine how niche and fitness differences are influenced by the presence of the parasitoid. We use a Bayesian approach to propagate uncertainty in model parameters through to niche and fitness differences, a method that should be widely applicable for understanding species coexistence processes in a range of systems. We show that parasitoids strongly influence the fitness differences between species and have inconsistent impacts on pairwise coexistence that would not be identifiable solely from reproductive rate – susceptibility relationships.

## Methods

### Study Species

We investigated a system of six naturally co-occurring *Drosophila* vinegar flies (*D. birchii* (BIR), *D. pallidifrons* (PAL), *D. pandora* (PAN), *D. pseudoananassae* (PSA), *D. simulans* (SIM), *and D. sulfurigaster* (SUL)) from a well-characterised Australian rainforest community (Hangartner et al., 2015; Jeffs et al., 2020; O’Brien et al., 2017), along with a co-occurring generalist parasitoid wasp from the genus *Trichopria* (Hymenoptera: Diapriidae, lab strain 66LD, SI 1). This parasitoid attacks *Drosophila* species at their immobilized pupal stage. Successful development of a parasitoid results in the death of its host. *Drosophila* can mount a variety of physiological defences to parasitoid attack, which can be costly (Fellowes & Godfray, 2000).

*Drosophila* species have long been used in experimental ecology since they are straightforward to culture under laboratory conditions, allowing high levels of replication and standardisation. The set of *Drosophila* species used here represents a significant fraction of the diversity of *Drosophila* species co-occurring at rainforest sites in northern Queensland, Australia (Jeffs *et al*. 2020). The six species span a range of body sizes (0.94 mg – 2.64 mg, mean wet mass of female adults) and phylogenetic divergences (SI 1, Table S1.1), have similar generation times, and can be distinguished visually (at least in the case of males). *Drosophila* are generalist feeders (Valadão et al., 2019) and in natural populations they experience high levels of parasitism, including from a suite of highly generalist natural enemies (Jeffs et al., 2020).

### Experimental design

We conducted a large, single-generation pairwise competition assay using a response-surface design following the guidelines of Hart *et al*. (2018), and in the presence or absence of pupal parasitoids. We used 25mm-diameter standard *Drosophila* vials (VWR International) containing 4 ml of cornflour-yeast-agar medium. This creates conditions where *Drosophila* compete for space to lay eggs, for larval food, and for pupation sites. Equal numbers of male and female *Drosophila* were introduced to each vial, so we refer to the number of founder pairs. Each two-species combination (15 combinations in total) was tested at different founding densities in a factorial design: (3 pairs of species A, 3 pairs of species B), (6A,3B), (9A,3B), (12A,3B), (3A,6B), (3A,9B) and (3A, 12B), with each combination replicated three times in each parasitism treatment. We also included monocultures of each species, with 12 replicates of a single founding pair, 6 replicates of 3, 9 or 12 pairs within each treatment, making a total of 990 experimental vials (360 single-species, 630 two-species). The entire experiment was split over three blocks, each with identical sets of combinations, but staggered by two days. The experiment was conducted in a single large incubator maintained at 26°C, with the location of trays within the incubator regularly rotated. Further details are given in SI 1.

### Initiation of treatments

Founding adults were offspring of adults drawn from mass-bred line cages and were raised in vials at moderate density (approximately 100 adults per vial) at a constant 23°C. These adults emerged 8-10 days before the start of the experiment, and so could be assumed to be sexually mature and mated. On the day preceding the start of the experiment the founding flies were lightly anesthetised under CO_2_ to be separated by sex. Males were added to the experimental vials immediately. Females were maintained at moderate density (∼50 adults per vial) in holding vials with fly medium and were transferred under light CO_2_ anaesthesia to experimental vials the following day, minimising the time interval between the addition of each species in multi-species assays. They were able to lay eggs and re-mate for 24 hours before being removed. Fig. 1 provides an overview of the experimental protocol.

**Figure 1.**
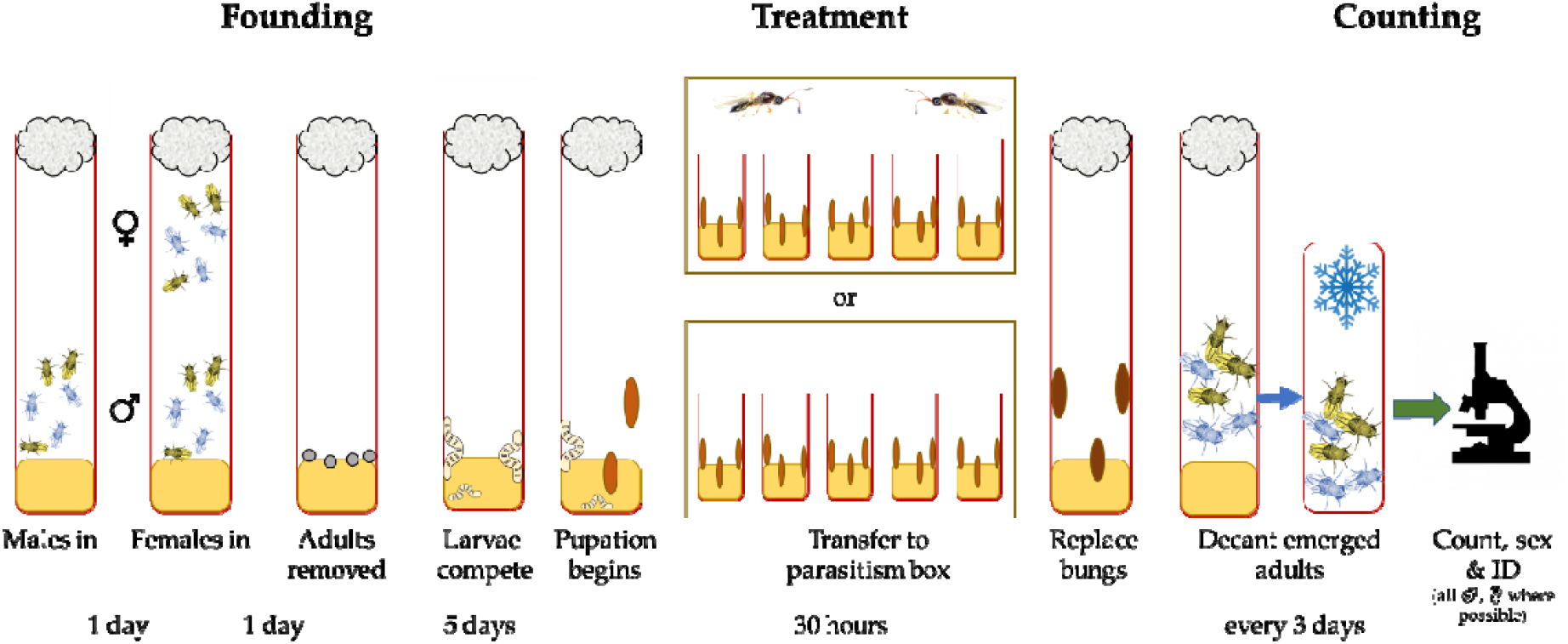
Summary of the experimental regime showing the founding process, treatment and data collection procedures described in the main text.

### Parasitism

At the onset of pupation, 5 days after the founder flies were removed, the fly vials were moved into Perspex parasitism boxes (35×20×14cm) containing up to 88 vials (SI 1). Vial positions within each box were randomised. The vial plugs were removed, and 50 female and 50 male mature and mated parasitoid wasps were added. This ratio of parasitoids to *Drosophila* was selected based on prior experience to result in a parasitism rate in line with field-parasitism levels of around 15% or greater (Jeffs et al 2020). The wasps were free to move between vials within each box. While it is possible that late-pupating *Drosophila* larvae could cross from one vial to another during this period, we only observed a handful of cases where 1-2 individuals of the ‘wrong’ species emerged from a vial, and very few pupae (<5 per box) were developed on the outside of the vials, suggesting that movement was negligible. After 30 hours, before the emergence of the fastest-developing *Drosophila* species, all wasps were removed with an aspirator, vial plugs were replaced, and the vials were returned to their original trays.

### Counting

We removed adult flies emerging from all vials every three days, minimising the risk that emerged adults started a subsequent generation. Flies in single-species vials were sexed and counted at each removal time, while flies in multi-species vials were sexed and identified at the end of the experiment (Fig. 1). There was a highly consistent 1:1 sex ratio across the single-species trials and in those 2-species trials where females of different species could be reliably distinguished (SI 2 Fig. S2.1). In the five of the combinations (BIR-PAN, BIR-PSA, BIR-SIM, PSA-PAN and PAL-SUL) where it was not possible to identify female flies to species we counted the males and inferred the total number of flies of each species assuming a 1:1 sex ratio.

## Modelling

### Determining growth rates

The larval competition of *Drosophila* often occurs within individual fruits that represent a discrete and temporary resource suitable for only a single generation of flies (Atkinson & Shorrocks, 1981; Shorrocks, 1991). We take as our underlying model of growth rates the Beverton-Holt model, without any carry-over between generations:

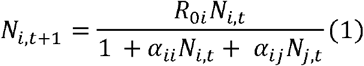

where *N*_*i,t*_and *N*_*i,l+t*_ are the size of the population of species *i* in the founding and emerging generations, respectively. *R*_*0i*_ is a species-specific generational reproduction term, *α*_*ii*_ represents the intra-specific competitive effect of species and *i* and *α* _*ij*_ are competition coefficients terms representing the effect of species *j* on *i*. It is important to emphasise that these *α* terms capture *all* impacts between species, including those through indirect means. The same function has been used for annual plants that have been the focus of most recent coexistence work (Bartomeus & Godoy, 2018; Levine & HilleRisLambers, 2009; Pérez-Ramos et al., 2019).

The mean number of adults emerging per founder was consistently lower in vials with a solitary founding pair compared to those with three founders (SI 2, Fig. S2.2). *Drosophila* use cues from other females when determining oviposition sites (Duménil et al., 2016; Wertheim et al., 2002), which could be causing this Allee effect. It is not possible to incorporate such positive density-dependence effects directly within the standard MCT framework without additional assumptions (Barabás et al., 2018; Schreiber et al., 2019). Since solitary founders are quite possibly a rather artificial scenario, we excluded data from all singleton pair trials from the model fitting.

### Inferring model parameters

The models were fit within a Bayesian framework and posterior distributions of the parameters were obtained from Hamiltonian Monte-Carlo sampling using STAN (Carpenter et al., 2017). We assumed a negative binomial error distribution and fitted separate overdispersion terms *ϕ* for each parasitism treatment. Since we counted both species, each two-species experiment contributed two data points. Although this contributes a small component of non-independence, the gain in total sample size is significant. Since we excluded the single-founder vials, our effective total ‘n’ across both treatments was 1476, to estimate a maximum of 86 free parameters in our most complex model (including overdispersion terms).

The competition α terms were each fit with an underlying Gaussian prior (mean=0, σ =1) and constrained to be positive, implying only competitive interspecific interactions, with no facilitation possible. Growth rate terms *R*_*0*_ were constrained to be positive, and fit with a weak Gaussian prior (mean=10, σ =10). Tests fitting α values with different constraints and priors did not give notably different results (SI 3).

We fitted and compared a sequence of candidate models of increasing simplicity. In the first, separate *R*_*0*_and terms were fitted for each treatment. In the second, a single set *α* of terms, but separate *R*_*0*_ terms were fitted across both treatments (i.e. parasitism affects emergence rate, but not competition coefficients). In the third model a single set of parameters were fitted across both treatments (i.e., parasitism has no effect). The three models were compared based on expected log pointwise predictive density (ELPD, a measure of out-of-sample predictive capacity), computed using the *loo* R package (Vehtari et al., 2017). This approach allows an assessment of whether there is sufficient evidence in the data to separate the parameters assigned to the treatment groups in a comparable manner to other information criteria such as AIC (Burnham & Anderson, 2002). All STAN model code and data to fit the models are available online. Posterior predictive checks and diagnostics are presented in SI 4.

### Pairwise coexistence criteria

The invasibility criterion for species coexistence requires each species to have a positive long-term low-density growth rate in the presence of the rest of the community (Chesson 2000). Following Godoy & Levine (2014), coexistence of a pair of species is determined to be possible, following Beverton-Holt model dynamics, when:

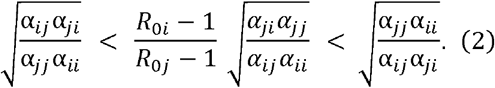

Equation (2) expresses the coexistence criterion as a balance between niche overlap (ρ) and fitness differences 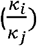 where:

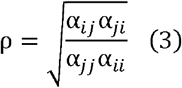

And

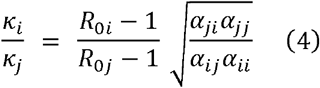

Following Chesson (2000) and Godoy & Levine (2014), we define the niche difference between two species as 1-ρ. Certain α values can result in a niche overlap ρ>1, implying a ‘negative niche difference’. This does not have a clear ecological meaning, and some previous authors have elected to define such cases as 0. However, since these negative1-*ρ* values can be ecologically interpreted as potentially identifying priority effect situations (Ke & Letten, 2018), we present the values as calculated. For both negative and zero values, coexistence is not expected under the MCT framework.

The *Drosophila* species we examined vary in their time to maturation. However, it is important to note that the mathematical expression of the invasibility criteria in a constant environment, equation (2), is not dependent on generation time. That is to say, the sole criterion for the invader species to expand its population while the resident is at an equilibrium is that its population size in its next generation will be larger, regardless of how long a generation takes.

For each pair of species, we calculated the niche and fitness differences across 3000 draws from the posterior distributions of parameter estimates. We partitioned the posterior to infer the support given by the data to each of the four outcomes of competition: coexistence, exclusion of either species or a situation of priority effects (see Fig. 3a). Note that the dynamics described by our models are fundamentally deterministic – these values express uncertainty in outcome given the data, rather than necessarily stochastic outcomes.

## Results

Based on model comparison, there was strong support for the expectation that parasitism reduces the growth rate of each species, but little support for differences in competition coefficients between parasitism and no-parasitism treatments. Our second model (with different *r* values for parasitism and no-parasitism treatments but common α values) performed best (Δ ELPD vs Model 1 = −43.02, vs Model 3= −34.03, SI4 Table S4.1).

The impact of the parasitoids on coexistence was inconsistent across species pairs (Fig. 2c). In three pairs of species the fitness difference was so large there was essentially no support for coexistence under either treatment. Of the remaining twelve pairs, five had higher support for coexistence under the parasitism treatment, while seven had less support (Table 1). In only two pairs (*D. sulfurigaster and D. pallidifrons, D. simulans* and *D. pallidifrons*) did the parasitism treatment cause the median estimate of fitness difference to cross the boundary delineating coexistence from competitive exclusion. In both cases the parasitism was associated with a moderate reduction in support for coexistence (Table 1). In the pair showing the largest change in support for coexistence between parasitism and no-parasitism treatments (+ 32 percentage points, *D. sulfurigaster* and *D. birchii*, Fig 2b), the median value did not cross the boundary. Strong positive correlations between fitted parameters, led by relationships between *R*_*0i*_ and α_*ii*_ terms, resulted in non-independence of estimates of the fitness and niche differences and non-eliptical posterior distributions (Fig. 2b,c).

**Table 1.**
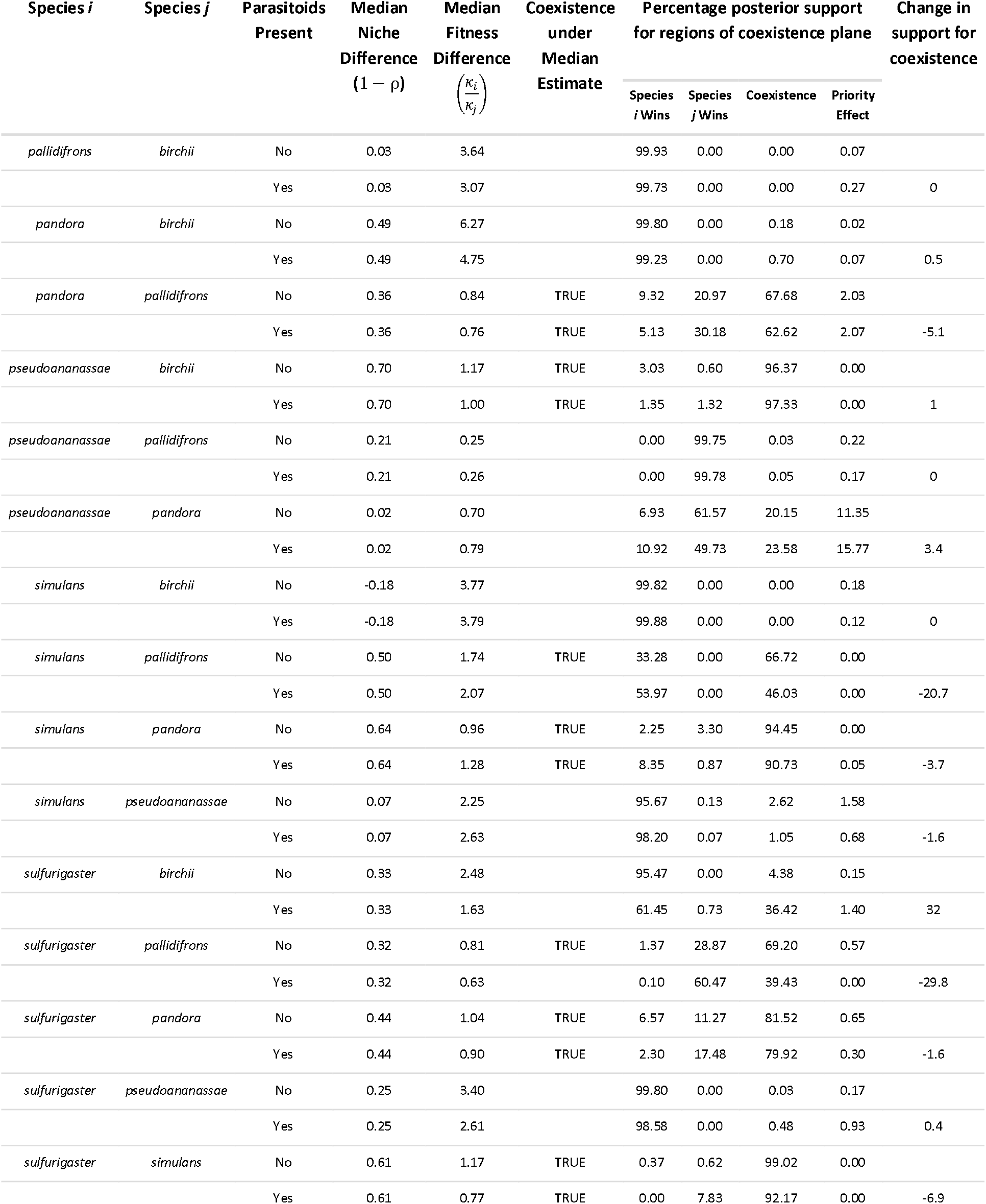
Summary of the posterior distribution across the coexistence plane between the two treatments for each species pair. The percentage of posterior draws that fall into the four major regions of the coexistence plane can provide a summary of the confidence in the outcome of a particular competition, given the data.

**Figure 2.**
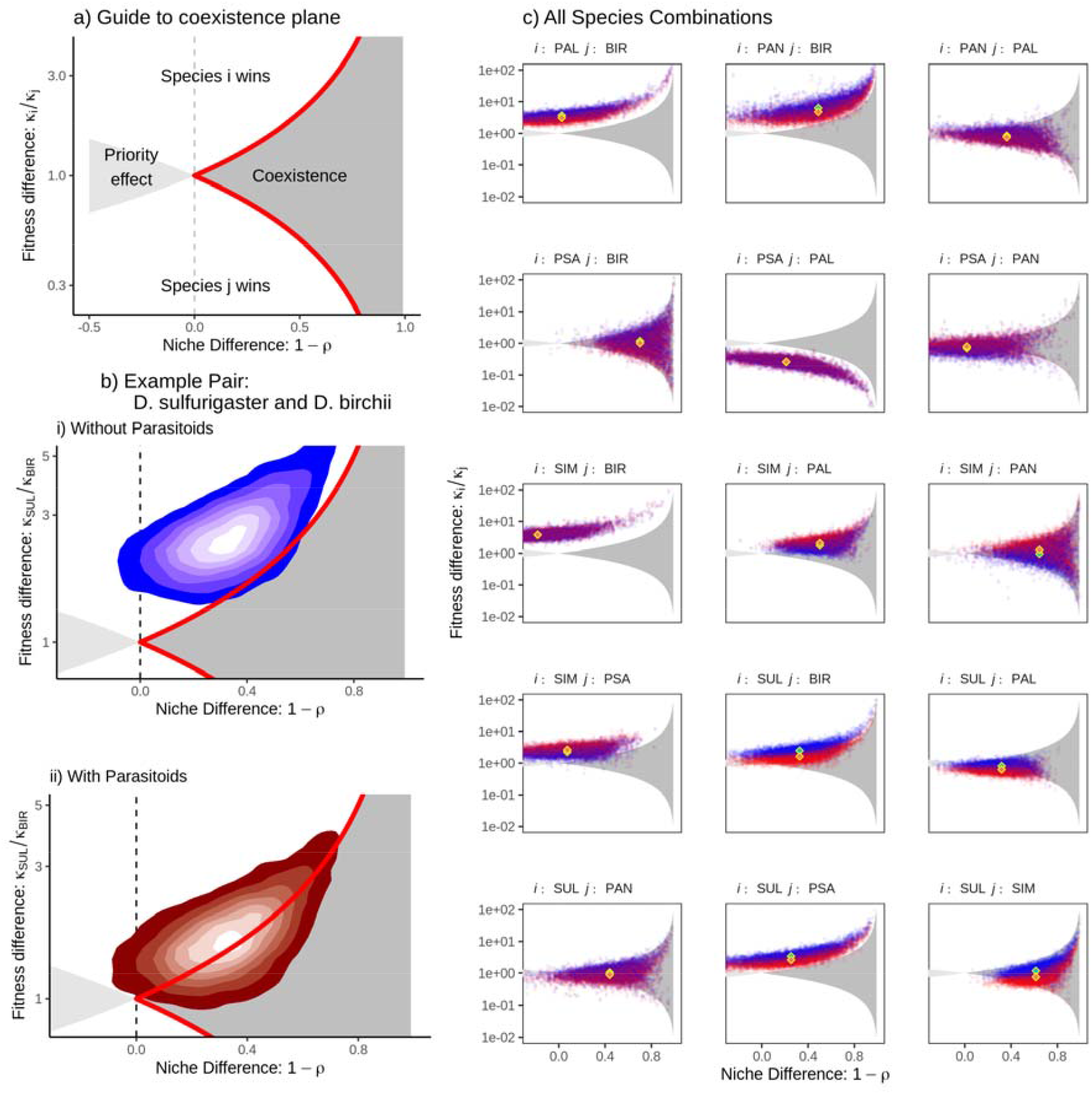
The likelihood of coexistence of *Drosophila* species pairs responds idiosyncratically to parasitism. a) Guide to interpretation of coexistence plane diagrams. The region of coexistence (dark grey, bordered by red line) defines the region where either species could invade from rare. The pale grey area denotes the region of ‘priority effects’, where neither species could invade the other. The white area is where fitness difference is large enough that the dominant species can competitively exclude the other species despite their niche difference. b) Full posterior distribution of niche and fitness differences of one example species pair (*D. sulfurigaster* and *D. birchii*) using the best-fitting model. Note the irregular, non-circular, shape ultimately derived from correlations between fitted vales in the posterior distribution and alpha terms contributing to both niche and fitness differences. c) Equivalent contour plots for all 15 species combinations. The median of the posterior distribution of parameters are shown with diamonds. Table 1 details the support assigned to each region for each pair under each treatment. Note that because the best model did not include varying competition coefficients () between treatments, only the fitness difference (y-axis) shifts between treatments.

The parasitism treatment reduced population growth rates, but we did not identify a reproduction rate – susceptibility trade-off across our six species (Fig. 3). The pairwise relationship between reproductive rate and susceptibility to parasitism (proportional reduction in *R*_*0*_) did not provide a useful heuristic in identifying the impact of parasitism on reducing fitness differences. Positive and negative reproduction rate – susceptibility relationships were both associated with increases and decreases in fitness differences.

**Figure 3.**
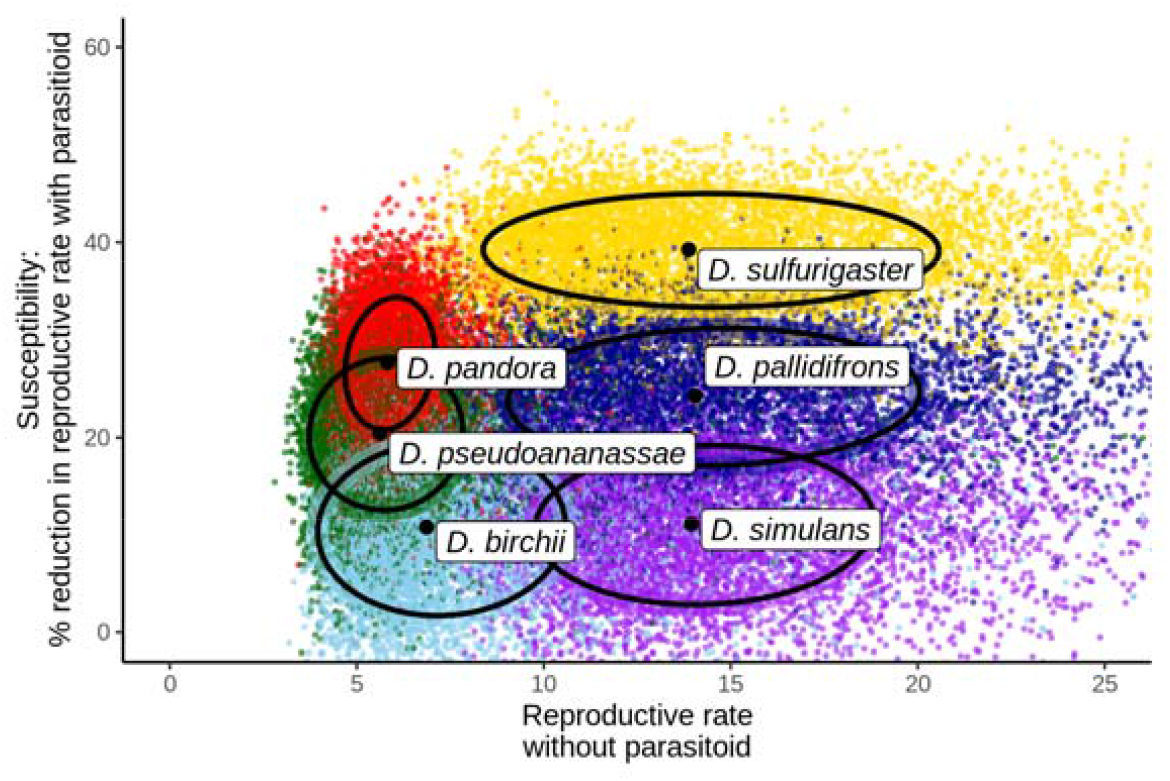
No clear reproductive rate – susceptibility trade-off across the six *Drosophila* species. Median value for each species (large black points) are plotted alongside a posterior distribution point cloud and fitted ellipse (60%, fitted assuming a multivariate-t distribution) to show the distribution of uncertainty for each species.

## Discussion

Our results demonstrate an inconsistent impact of natural enemies on the pairwise coexistence of a guild of competing species. *Drosophila* species showed differential susceptibility to parasitism, which manifests in shifts of the fitness differences in the coexistence plane. Natural enemies shift the fitness difference in favour of the species that is less susceptible, but this does not necessarily enhance coexistence. Across our six species, no general reproductive rate – susceptibility trade-off was observable and the presence of parasitoids did not lead to a consistent shift towards (or away from) a state of potential coexistence. At the pairwise level, reproductive rate – susceptibility relationships did not provide a useful guide to the impact of parasitoids on coexistence. This is in line with results of Petry et al (2018) where seed predators reshuffled the results of competition and emphasises the insufficiency of fecundity as a substitute for competitive ability in practice. However, in contrast to Petry et al (2018) we did not observe any cases of ‘overshooting’ the zone of coexistence and reversing the victor of competition.

Fully quantifying pairwise competitive ability requires considerable effort (Hart et al., 2018), that scales quadratically with the number of species to be assessed. Since an increased fecundity increases competitive ability, taking shortcuts based on direct measured of fecundity and susceptibility to determine the role of natural enemies is appealing. However, our results provide further evidence that this represents a dangerous trap, and there is no substitute to full characterisation of interspecific competition. Predation can only favour coexistence when the more susceptible species is otherwise competitively dominant, not just more fecund. This underscores the importance and challenges of accurately measuring competition terms (α) to understand the consequences of any given factor’s influence on coexistence.

### Impact of enemies on competition coefficients

In natural systems, both direct effects of mortality and impacts on the interaction between species will determine the impact of consumers on species coexistence (Pringle et al., 2019). Our study design allowed parasitoids to travel between vials, opening the prospect of either associational susceptibility or resistance mediated through parasitoid foraging behaviour (Abrams, 2010; Holt & Kotler, 1987). If these competitive interaction modification effects were strong enough to be detectable, they would manifest as shifts in the fitted α values and change the niche differences between parasitism and no-parasitism treatments. However, our model selection procedure suggested that modelling changes to the competition coefficients matrix between treatments leads to overfitting i.e. there were no identifiable impact of parasitism on pairwise competition coefficients. Resolving the relative importance of direct effects of predators versus their indirect effects on other species is a long standing challenge (Werner & Peacor, 2003). Our study suggests that in this case the direct effects are dominant. However, mechanisms that can be modelled as shifts in single-species parameters (e.g. *R*_*0*_) are inherently more identifiable statistically than changes to multi-species processes (e.g. α _*ij*_) as they can be determined more directly from simpler experiments.

Nevertheless, our model selection test is a rather blunt all-or-nothing approach. It may well be that only particular competition coefficients are strongly affected by the parasitoid. Indeed, in our Model 1, while most of the between-treatment α differences cluster tightly together, several show large divergences (SI 4 Fig. S4.14). There is no clear predictor for the terms that show these large shifts, occurring between different species. A more complex model-fitting approach incorporating regularisation could potentially determine whether such cases are identifiable effects rather than model-fitting noise.

Our single-generation design meant that density-mediated apparent competition derived from an increased total population of natural enemies was not directly detectable. If we had run a multi-generational experiment, the results could be expected to be complex: Sommers & Chesson (2019) show that the relative balance between apparent and resource competition is influenced by prey defensive actions. Methods to compare resource competition and predation effects directly in parameterised models are being developed (Shoemaker et al., 2020) and will be an exciting area for future research.

### Model fitting considerations within coexistence theory

Recent meta-analyses have emphasised the importance of parameterised models for understanding mechanisms of coexistence (Broekman et al., 2019), particularly since large-scale global environmental changes will create novel communities (Alexander et al., 2016). Modern coexistence theory defines a sharp line between coexistence and the inability to invade from rare. This mathematical clarity and precision represents a major advantage, but poses challenges to the incorporation of inevitable experimental uncertainty, even before comparison to field contexts. It is important to reiterate that the uncertainty of coexistence we discuss should be interpreted as statements of uncertainty in the underlying data, rather than implying a probabilistic outcome. The precision of estimates of parameters can differ between treatments, which can have the effect of changing the support for coexistence, even if the central estimate does not change. Furthermore, these values should not be directly interpreted as some measure of ‘distance from coexistence’, although the concepts may be related. The framework of structural stability is better equipped to address such questions (Godoy et al., 2018; Saavedra et al., 2017).

Nonetheless, parameter fitting can have high data demands, and key ecological model parameters are often not fully separable (Bolker, 2008). For example, in a simple model of competition similar predictions could be made by lowering the growth rate term and increasing the competition term. With a well-designed experiment and sufficient data, uncertainty can be minimised, but in any realistically-sized competition experiment, separability of parameters can be a significant but largely underacknowledged issue. This problem is particularly pertinent when working within the framework of modern coexistence theory, where conclusions are drawn from comparisons between ratios composed of multiple parameters from the underlying model (Barabás et al., 2018; Song et al., 2019). Failing to address these issues could dramatically affect the confidence with which conclusions are drawn from experimental data.

Understanding how the interrelationships between parameters affect uncertainty in the derived quantities based upon them is difficult a priori, since the contribution of different correlations will be contingent on both the underlying ecology and the experimental design. In the context of modern coexistence theory, even a moderate confounding of underlying growth rates and intra-specific competition terms may have a significant effect on the conditions for coexistence. In our study, the posteriors were very often curved - posterior draws generating large niche differences were associated with larger relative fitness differences. This has the beneficial effect of often increasing the confidence with which pairs of species are located within regions of the coexistence plane. Although our individual estimates of model parameters may be rather uncertain, we can often be comparatively confident in the result of competition.

This effect can be attributed to indeterminacy, and hence correlation in the posterior, between growth rates and intra-specific competition terms. One consequence of this can be seen clearly when the condition for coexistence (equation 2) is re-arranged in a simplified form as:

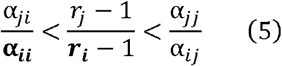

Where (as in our data) there are strong positive correlations in the posterior draws of α_*ii*_ and *r* _*i*_ (shown in bold), this would tend to affect both sides of the inequality simultaneously, increasing the tendency that it maintains its value.

Fitting parameterising models with ecological data is almost always both challenging and illuminating in equal measure. Because the fitted values of the parameters inevitably correlate to some degree, propagating parameter uncertainty from a multi-parameter posterior distribution gives a better understanding of the uncertainty associated with the key quantities of modern coexistence theory (Mordecai, 2013; Uricchio et al., 2019). Multi-dimensional posteriors can pose considerable challenges both to intuition and to visualise. However, approaches that directly propagate standard errors associated with individual parameters (e.g. Matías et al. 2018) or estimate uncertainty by bootstrapping observations (e.g. García-Callejas et al. (2020) are unable to capture these effects, and risk misrepresenting the confidence that can be placed in results. Although we took an explicitly Bayesian approach using weak priors, similar methods could equally be followed using multi-dimensional likelihood surfaces computed with a frequentist approach.

### Limitations of simple laboratory experiments

The modern coexistence theory approach is agnostic with respect to the particular factors involved but will always be limited by the scope of the empirical data available. Experiments such as this one can rarely replicate full natural lifecycles of animals or capture the full range of conditions that occur throughout their habitat (Hart et al., 2018). Impacts on one life stage can mitigate differences at another (Moll & Brown, 2008) and our method does not capture differences in reproductive lifespan. Equally, our experiment uses a highly simplified representation of the consumer pressure experienced by natural communities of *Drosophila*. We included a single pupal parasitoid species; in their natural habitats these species exist in a complex trophic network (Jeffs et al., 2020) including specialist parasitoids and larval parasitoids (which may directly influence larval competitive performance).

Coexistence may only be possible at particular spatial scales (Hart et al., 2017), or through spatial mechanisms not reflected in our experiment. The laboratory experiments cannot capture environmental heterogeneities hypothesised to aid coexistence (Chesson, 2000; Kuang & Chesson, 2010). Despite these limitations, we believe that our experiment is still instructive in isolating particular aspects for further investigation.

The *Drosophila* species we investigated co-occur in their natural habitat of Australian rainforest (Jeffs et al., 2020), although it is important to note that this is not necessarily evidence that they can coexist in the sense of mutual invasibility (Siepielski & McPeek, 2010). Furthermore, pairwise coexistence is not the same as community-level coexistence, and coexistence or otherwise between pairs does not necessarily translate to the multispecies community (Song et al., 2019), although they are likely to be at least somewhat related (Broekman et al., 2019; Friedman et al., 2017). A diversity of mechanisms can allow multispecies coexistence where pairwise coexistence is not possible (Letten & Stouffer, 2019; Levine et al., 2017; McPeek, 2019).

## Conclusion

Our experiment demonstrates that natural enemies can change the result of competition between species pairs, but that this effect is not consistent or necessarily predictable from pairwise reproduction rate - susceptibility relationships, reiterating the value of analysing parametrised models (Siepielski & McPeek, 2010). Further work will be needed to investigate whether trait differences impact the niche and fitness differences observed in our system, and more experimental studies of the impact of natural enemies on coexistence are needed to form the basis for a more general understanding of their impacts beyond plants (Viola et al., 2010). Modern coexistence theory is emerging as a powerful tool to investigate a range of biotic and abiotic impacts on coexistence experimentally (Lanuza et al., 2018; Matías et al., 2018). However, since the precision with which growth rates and competition coefficients can be estimated is limited (even within large studies such as ours), the fundamental inseparability of niche and fitness differences can impact the likelihood assigned to coexistence between any given species pair. Although these challenges are significant, we hope our approach demonstrates that they can be readily and transparently addressed.

## Acknowledgements

We thank the Hrček lab and the Oxford Fly group for advice and use of facilities, Chia-Hua Lue for advice on parasitoid taxonomy, František Sládeček and Jamal Holle for assistance maintaining cultures, and Mark Wong for constructive comments on the manuscript.

## Notes

**Funding:** The work was supported by NERC grant NE/N010221/1 to OTL and a grant from the China Scholarship Council to JC. JCDT was also funded through NERC grant NE/T003510/1

https://www.github.com/jcdterry/TCL_DrosMCT

### Competing Interest Statement

The authors have declared no competing interest.

### Summary of Updates

This revised version uses a slightly different core model and places different emphasis between the results.

