## Supplementary material for "Natural enemies have inconsistent impacts on the coexistence of competing species": SI 1-4

### SI 1 - Supplementary Methods

Chris Terry, Jinlin Chen & Owen Lewis

#### Species details

Prior to the start of the experiment, all isofemale *Drosophila* lines had been maintained in laboratory monocultures for 20-40 generations. Mass bred lines were constructed from isofemale lines and were maintained in large population cages for half a year. Drosophila medium contains 8% corn flour, 4% yeast, 5% sugar, 1% agar, and 1.67% methyl-4-hydroxybenzoate (concentration is measured by weight/volume of water).

| Species | Code | Subgroup | Distribution | Wet Weight (mg) |
|---|---|---|---|---|
| *D. birchii* | BIR | montium | highland-biased | 0.94 |
| *D. pallidifrons* | PAL | nasuta | highland-biased | 2.24 |
| *D. pandora* | PAN | ananassae | lowland-biased | 1.14 |
| *D. pseudoananassae* | PSA | ananassae | lowland-biased | 1.01 |
| *D. simulans* | SIM | melanogaster | only found at one lowland study site | 1.30 |
| *D. sulfurigaster* | SUL | nasuta | widespread | 2.64 |

**Table SI 1.** Details of *Drosophila* species used in laboratory experiment. Although all the species co-occur within their Australian rainforest habitat, different species are more abundant at particular elevations.

The parasitoid wasp used is as-yet unnamed, but belongs to the *Trichopria* genus (Hymenoptera: Diapriidae). Its Smithsonian National Museum of Natural History (NMNH) reference vouchers are USNMENT01557121, USNMENT01557273 and USNMENT01557254. The parasitoid has been maintained in laboratory monocultures (as lab strain 66LD) for 2 years after collection from the field from the same community as the hosts. During culturing, *Drosophila melanogaster* was used as a host species to reduce specific selection between the parasitoid and the experimental *Drosophila* species used in the laboratory.

#### Images of experimental set-up.

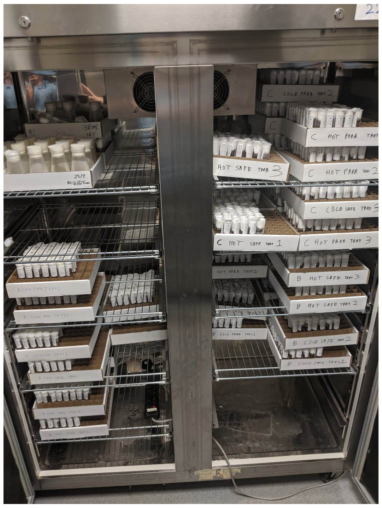

**Figure S1.1** Individually plugged vials in which founder adults laid eggs and larvae competed. Vials were mainainted in an incubator at constant temperature.

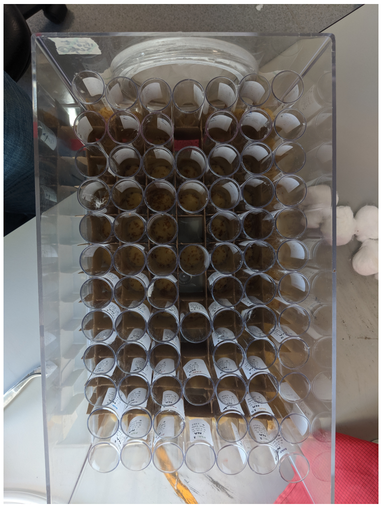

**Figure S1.2** Vertical view of unplugged vials within parasitism boxes. Empty vials containing parasitic wasps were placed in the gaps and the wasps could freely move between the tubes within each box.

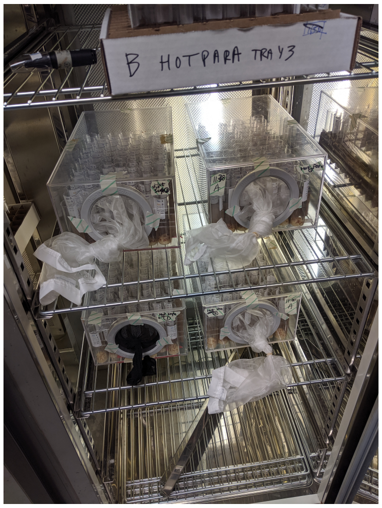


**Figure S1.3** Parasitism boxes (with lids in place) were maintained in the incubator during the parasitism period.

#### Location Randomisation Procedure

Within the incubator, vials were arranged in a randomised, but structured arrangement (Fig. S1.4 and S1.5). The structure was necessary to facilitate handling. In all cases the principle that each vial had an equal change of being located at any point was maintained. Within each treatment, within each block, there was one tray for single-species assays, and two for multi-species assays. Species combinations were kept in rows, but the order of the numbers of each species used was randomised.

Unfortunately, a mistake lead to parasitoid wasps being released into the a wrong box. Block C, Parasitoid(+) box number 2 did not receive parasitoids as planned, while Block C, Parasitoid(-) box number 1, did get parasitoids when it should not have done. This was corrected by reassigning vials to treatments, but had the result that certain combinations had one greater or fewer data point per treatment than was planned.

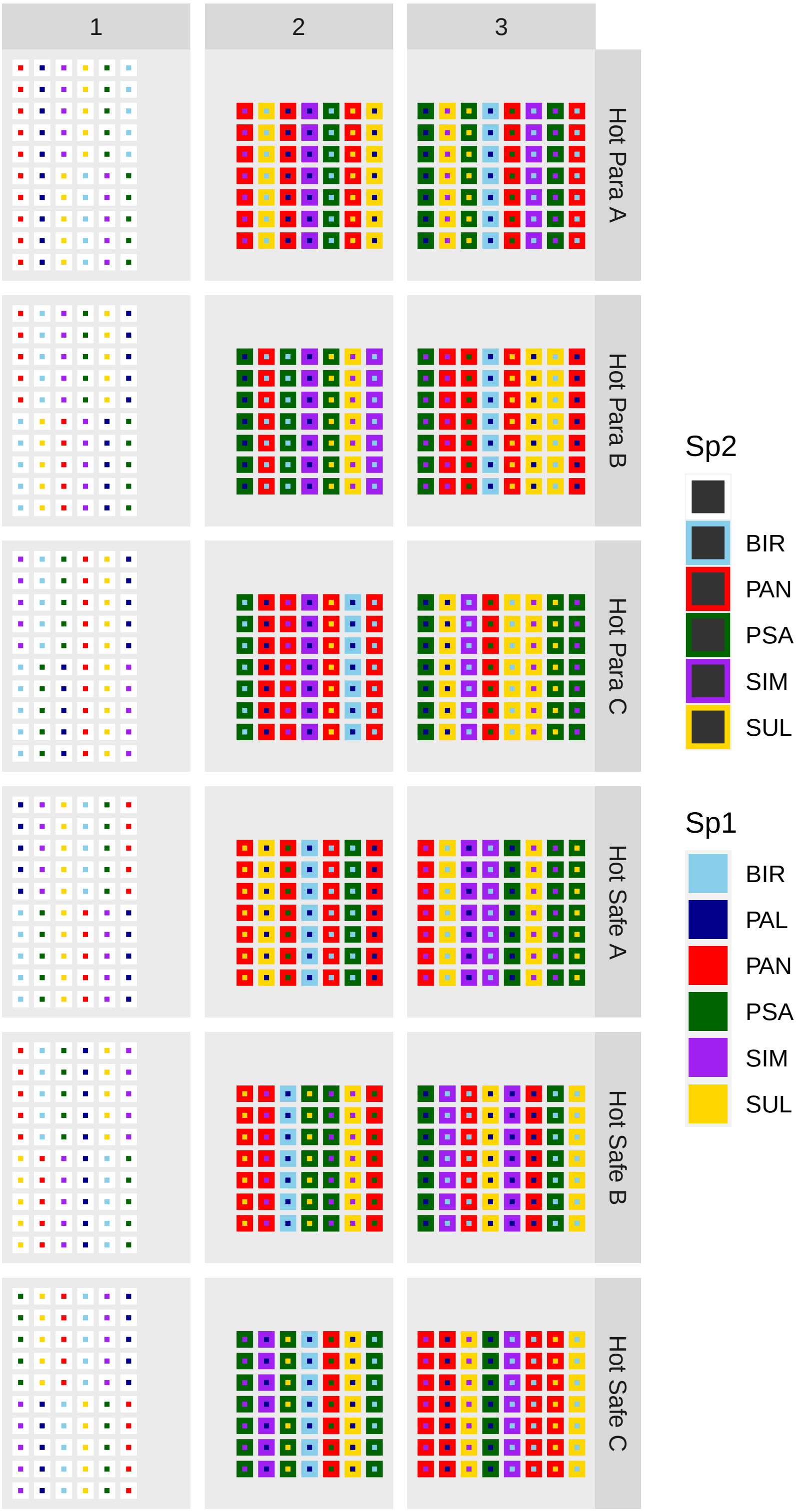


**Figure S1.4** Randomisation of vials in trays by species combination for founding and larvae competition. Species combinations were maintained in lines to facilitate rapid handling. Within each sub-block, the order of the founder density levels was randomised. The distinction between species '1' and '2' is arbitrary.

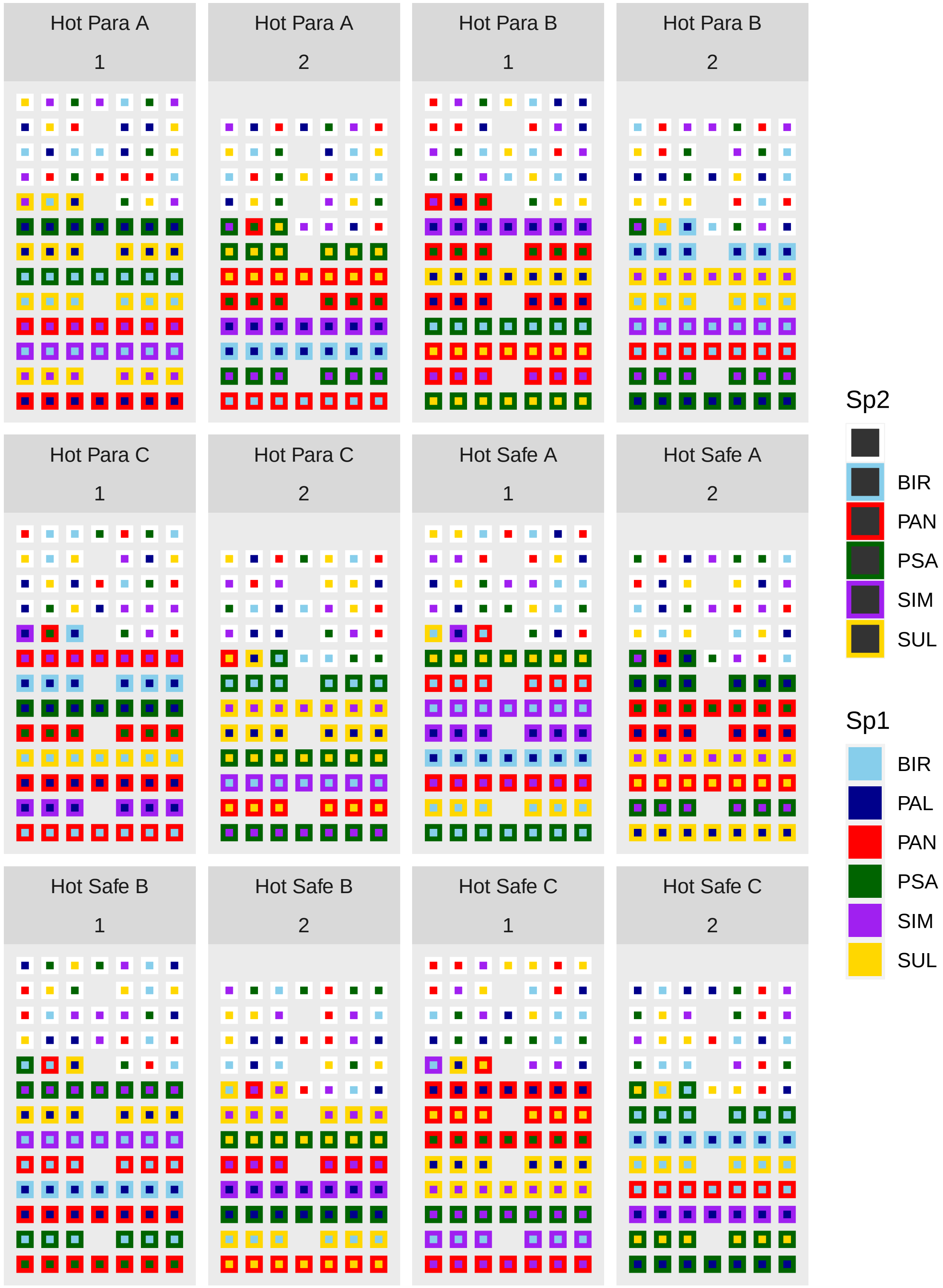


**Figure S1.5** Randomisation of vials into parasitoid treatment boxes. For the multi-species vials, the order of the founder density levels was randomised within each line. The distinction between species '1' and '2' is arbitrary.

#### Temperature Treatment

Our initial experimental design included a low-temperature treatment (20°C). This followed identical procedures, although some incubation times were lengthened to account for slower development times. However, too few adult flies emerged in the 20°C treatment for reasonable identification of competition parameters and we do not consider this cold-temperature treatment further.

### SI 2 - Model Design - Allee Effect & Sex Ratio

Chris Terry, Jinlin Chen & Owen Lewis

#### Sex Ratio

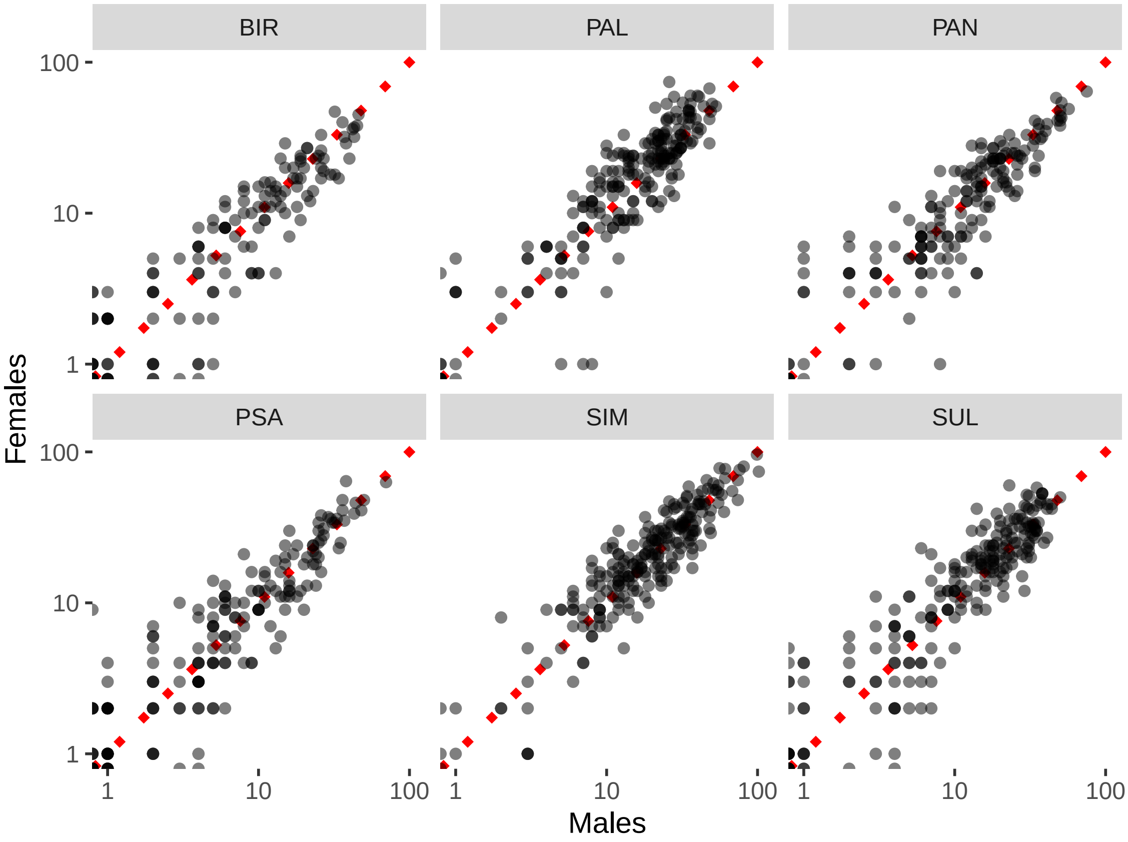


**Figure S2.1.** Consistent sex-ratio of emergent adults, faceted by species. Across all six species, a binomial GLM estimated only a very slight bias ($\beta$ coefficient = 0.065) towards females.

#### Allee Effects

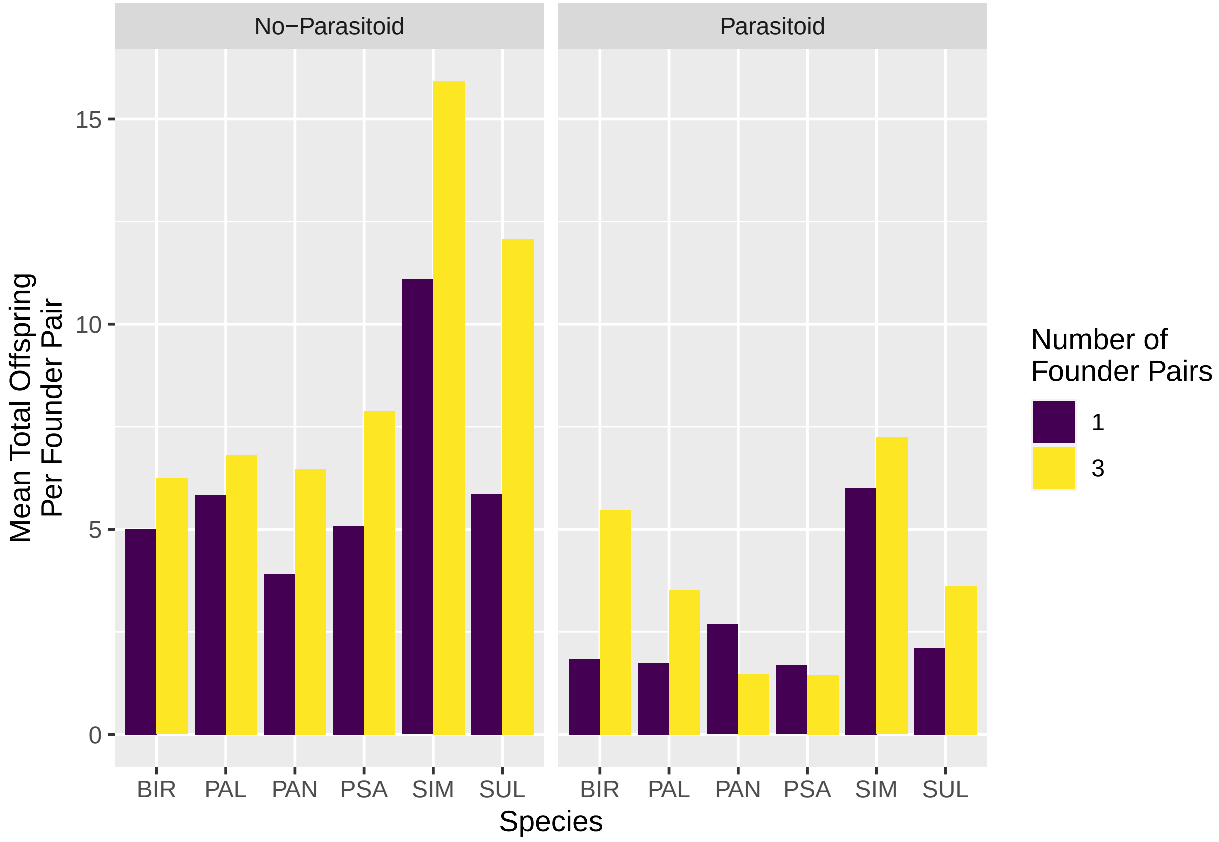


**Figure S2.2.** Per-female offspring is consistently lower for singleton founder pairs than triplets in the monoculture trials.

### SI 3 - Assessing Sensitivity to Priors

Chris Terry, Jinlin Chen & Owen Lewis

In our model, the $r$ parameters were relatively straightforward to estimate, and there is good reason to be confident that the choice of prior is not particularly influential. However, for the competitive coefficents ($\alpha$) the fits were considerably more uncertain, and there is a greater potential for the choice of prior to be influential. In the main text, we present results where the prior on each $\alpha$ parameter is constrained to be positive (i.e. exclude the possibility for facilitation) and conditioned on a zero-centered Gaussian distribution with standard deviation 1. To confirm that the choice of prior is not having an outsized effect, we tested tighter ($\sigma = 0.1$) and looser ($\sigma = 10$) priors on the $\alpha$ values, as well as a test where the lower $\alpha$ limit was reduced from 0 to -0.025 (lower values could lead to negative growth rates which are incompatible with the negative binomial error distribution).

Only the very tight prior resulted in markedly different $\alpha$ posterior distributions, indicating that our results are not overly influences by our choice of priors (Fig S3.1). Only one interaction coefficient had a posterior that notably included facilitatory values ($\alpha_{PAN,BIR}$). We have no particular explanation about why this interaction is negative, and can only assume it derives from data variability. As it is only a single term, we consider it unlikely to have a large influence on our overall results.

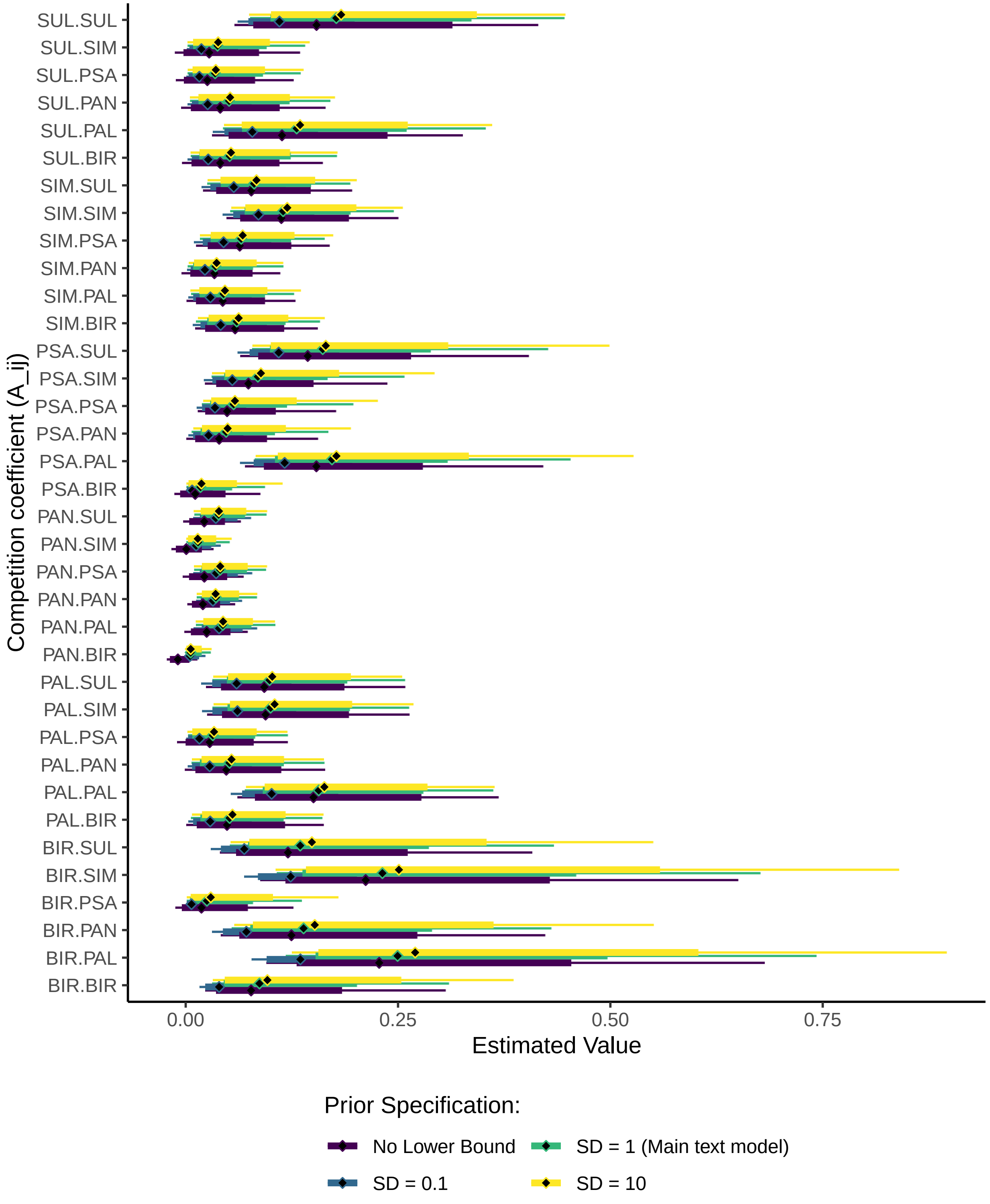


**Figure S3.1** Fitted posterior distributions of competitive coefficient terms under different prior assumptions.

### SI 4 - Additional Results

Chris Terry, Jinlin Chen & Owen Lewis

This appendix details the results of additional analyses to support the results in the main text.

A. Full results from model comparison

B. Median estimates for model parameters

C. Full posterior distribution of model parameters

D. Diagnostic plots for the model fitting, including posterior predictive plots

E. Results from maximal model including predation effects on competitive terms

#### A. Model comparison

| | Δ ELPD | Standard Error of Difference | Effective Number of Parameters | Standard Error |
|---|---|---|---|---|
| model2 | 0.0000 | 0.0000 | 48.5480 | 2.7322 |
| model3 | -34.0281 | 8.1397 | 43.6685 | 2.6892 |
| model1 | -43.0215 | 10.1924 | 76.6309 | 5.2007 |

**Table S4.1** Results from model comparison through expected log-pointwise predictive density (ELPD), computed using Pareto-smoothed importance sampling with the loo R package (Vehtari *et al* 2019). Models differ in the degree of divergence in fitted parameters between parasitism treatments. Model 1: separate $\alpha$ and $r$ values (representing the hypothesis that parasitism affects both growth rate and competition terms between species.) Model 2: joint $\alpha$ values, but variable $r$ (the hypothesis that parasitism affects growth rates, but not competition terms), Model 3: joint $\alpha$ and $r$ values (the hypothesis that parasitism has no effect). Differences in ELPD suggest strongest support for Model 2.

#### B. Median Parameter Values

Despite a major point of the paper being that the full posterior is essential to fully understand the uncertainties in parameter estimates, we appreciate that for some purposes a single point estimate is useful. Table S4.2 and Table S4.3 therefore present median estimates for the key model parameters.

| Species | Parasitism | Safe |
|---|---|---|
| BIR | 6.135 | 6.865 |
| PAL | 10.647 | 14.053 |
| PAN | 4.198 | 5.822 |
| PSA | 4.467 | 5.612 |
| SIM | 12.396 | 13.951 |
| SUL | 8.419 | 13.875 |

**Table S4.2** Median values for growth rates ($r$) in the two treatments

| | BIR | PAL | PAN | PSA | SIM | SUL |
|---|---|---|---|---|---|---|
| BIR | 0.0869 | 0.2495 | 0.1388 | 0.0250 | 0.2316 | 0.1348 |
| PAL | 0.0517 | 0.1566 | 0.0516 | 0.0319 | 0.1000 | 0.0987 |
| PAN | 0.0058 | 0.0437 | 0.0356 | 0.0406 | 0.0146 | 0.0388 |
| PSA | 0.0175 | 0.1723 | 0.0477 | 0.0564 | 0.0850 | 0.1613 |
| SIM | 0.0601 | 0.0446 | 0.0352 | 0.0654 | 0.1147 | 0.0806 |
| SUL | 0.0518 | 0.1307 | 0.0511 | 0.0346 | 0.0375 | 0.1773 |

**Table S4.3** Median values for competition coefficients ($\alpha$) laid out in standard matrix form ($\alpha_{row,col}$).

#### C. Parameter posterior distributions (selected model)

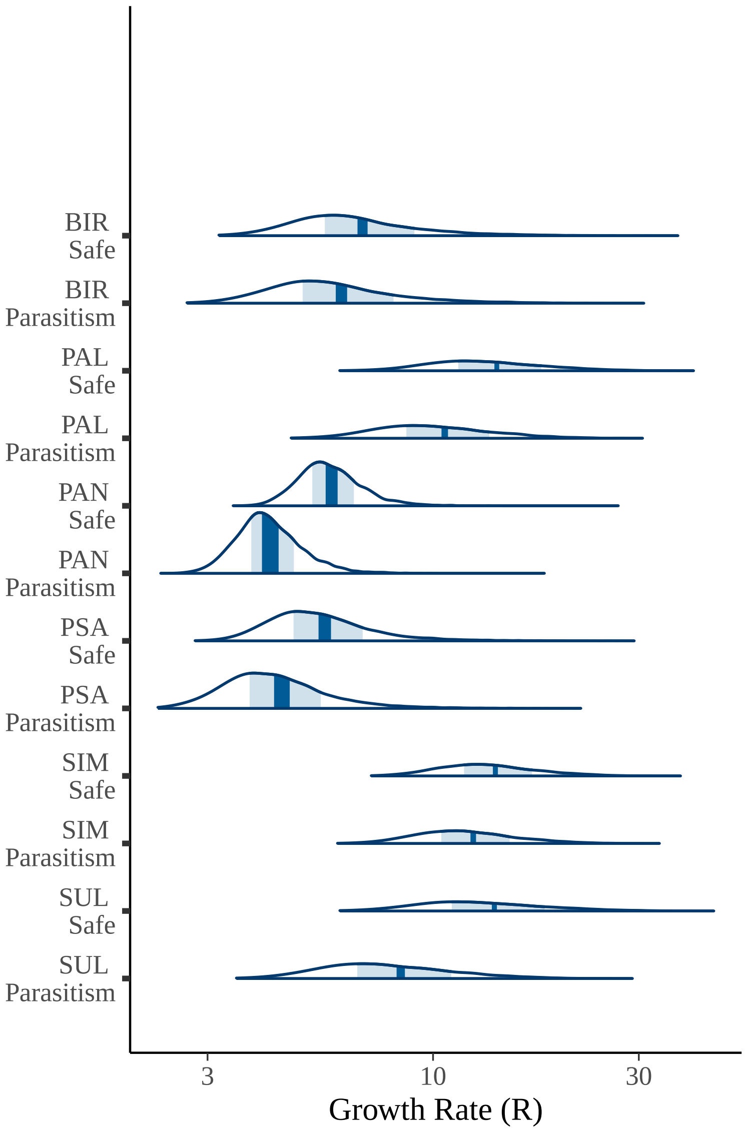


**Figure S4.1** Posterior distribution of growth rates under the two treatments. Shaded area shows 50% inner interval. N.B. log-scale.

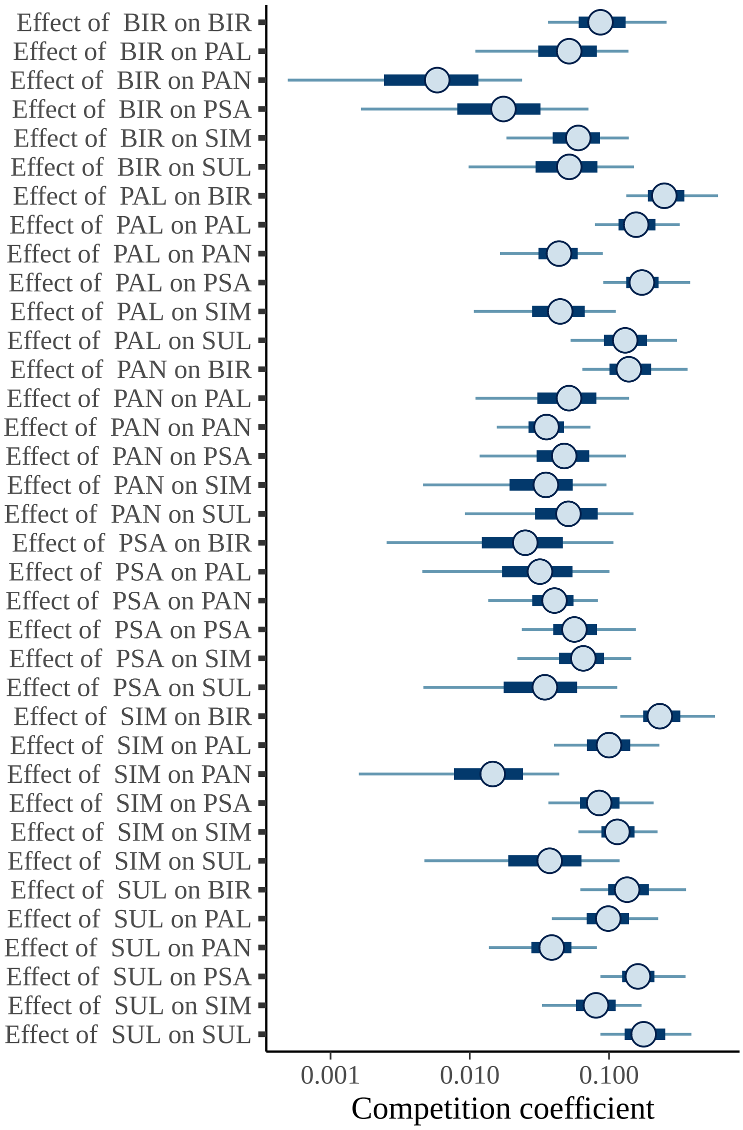


**Figure S4.2** Posterior distribution of competition parameters. Wide bar shows 50% inner interval, line shows 90% interval, circle shows median value. N.B. log-scale.

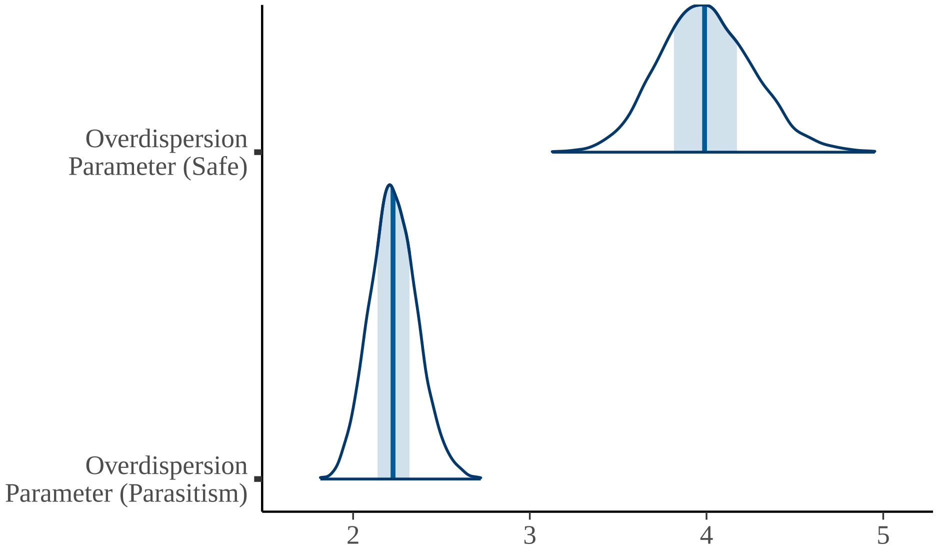


**Figure S4.3** Posterior distribution of the $\phi$ parameters, where the shaded area shows 50% inner interval. Note that in the parameterisation we used (`neg_binomial_2()`), the variance is defined as $\mu + \frac{\mu^2}{\phi}$. Therefore the inverse of $\phi$ controls the overdispersion, which is scaled by the mean value. $\phi$ is lower in the parasitoid treatment, suggesting that parasitism increases the variance in the next generation.

#### Relative uncertainty in components of fitness differences

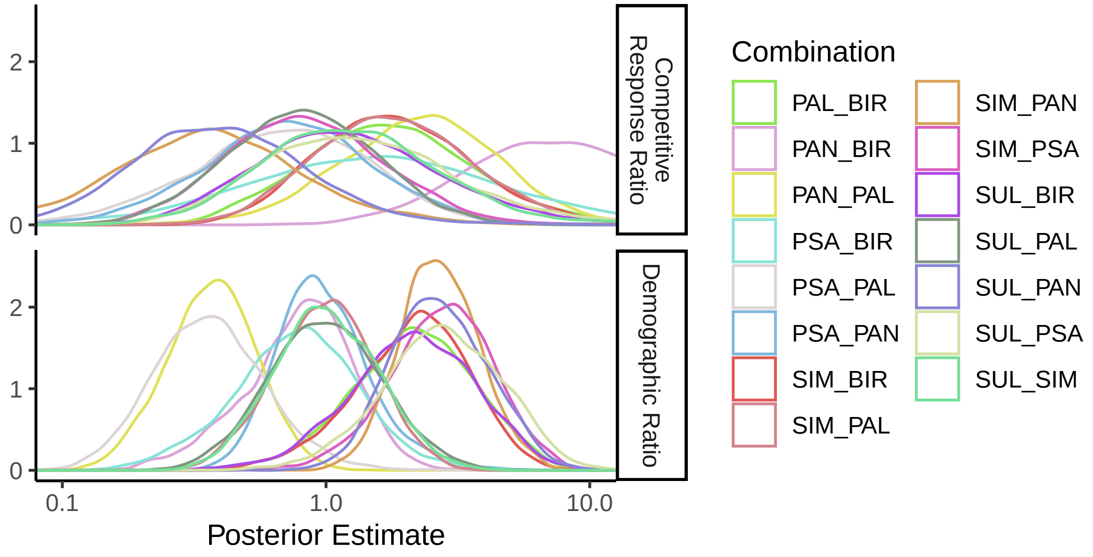


**Figure S4.4** Posterior distribution of the two components of the fitness difference. The uncertainty in the 'competitive response ratio' $\sqrt{\frac{\alpha_{ij}\alpha_{ii}}{\alpha_{ji}\alpha_{jj}}}$ is much larger than the demographic ratio $\frac{r_j - 1}{r_i - 1}$ (mean coefficient of variation 32x larger).

### D. Model Diagnostics

#### Overall Predictive Capacity

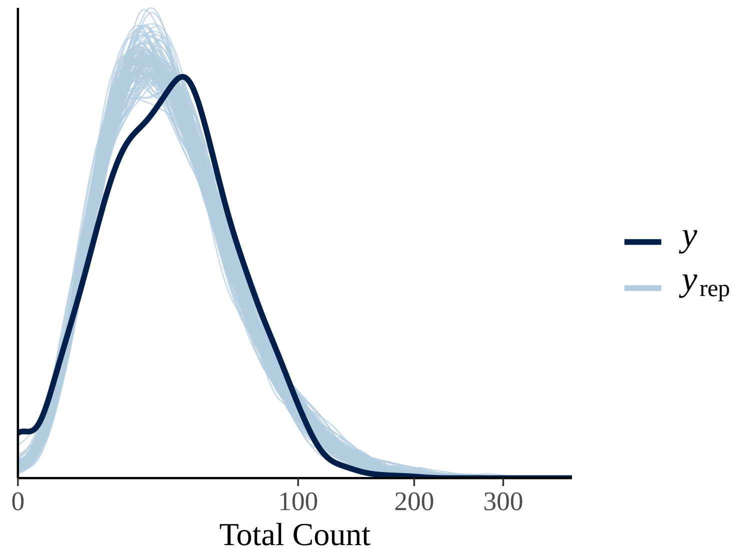


**Figure S4.5** Overall predictive capacity of best fit model. 100 sets of predictions from the posterior (blue), are compared to observed values (black). It can be seen that the model captures the overall spread of the data well, but does not pick up the low-growth rate observations (i.e. those cases where no adults were observed to emerge). N.B. the square-root X-scaling to magnify small values.

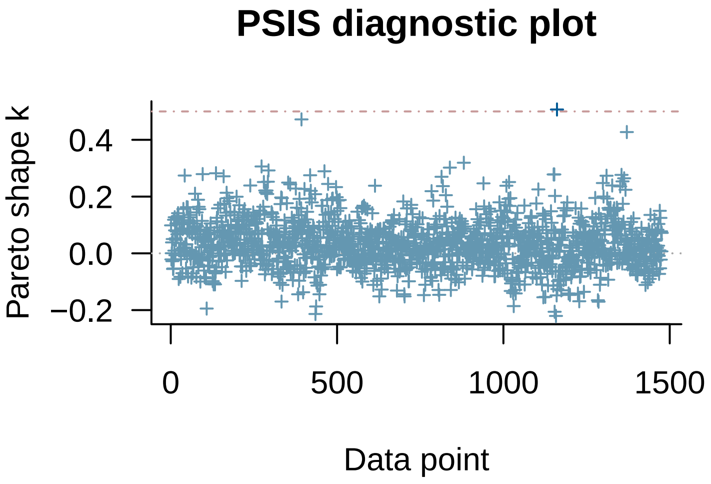


**Figure S4.6** Test for model miss-specification, using Pareto-smoothed Importance Sampling to test for strongly outlying data points. No points exceeded the threshold of 0.7 (Gabry *et al.* 2019).

#### Species-level Predictive Capacity

Figures S4.7 to S4.12 present predictions of the selected model against the raw observations. Because of the multi-dimensional nature of the data, the observations are partitioned by parasitoid treatment, and by the type of competition.

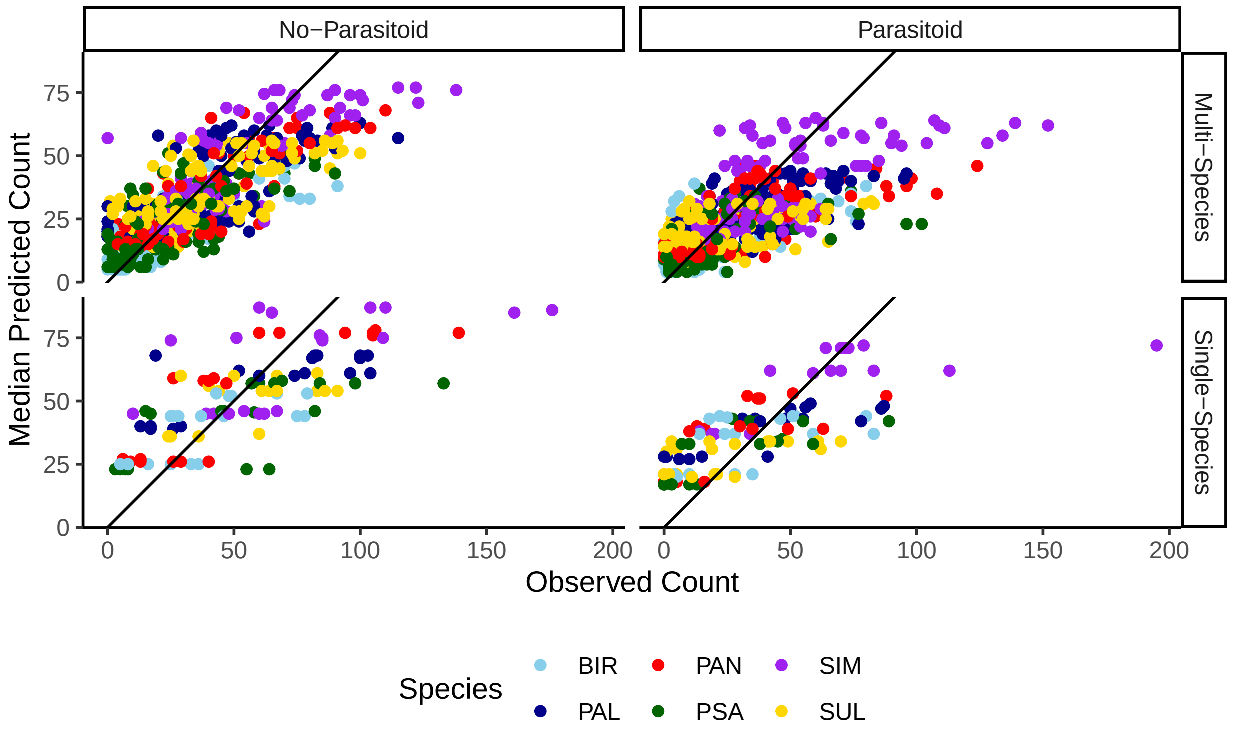


**FigS4.7** Overall model predictive accuracy using median posterior values. Plot is faceted by treatment and whether the count was from a single-species or a multi-species trial. Focal species are coloured. In general the model performs well given the inevitable variation, but struggles to capture outlier values across all species.

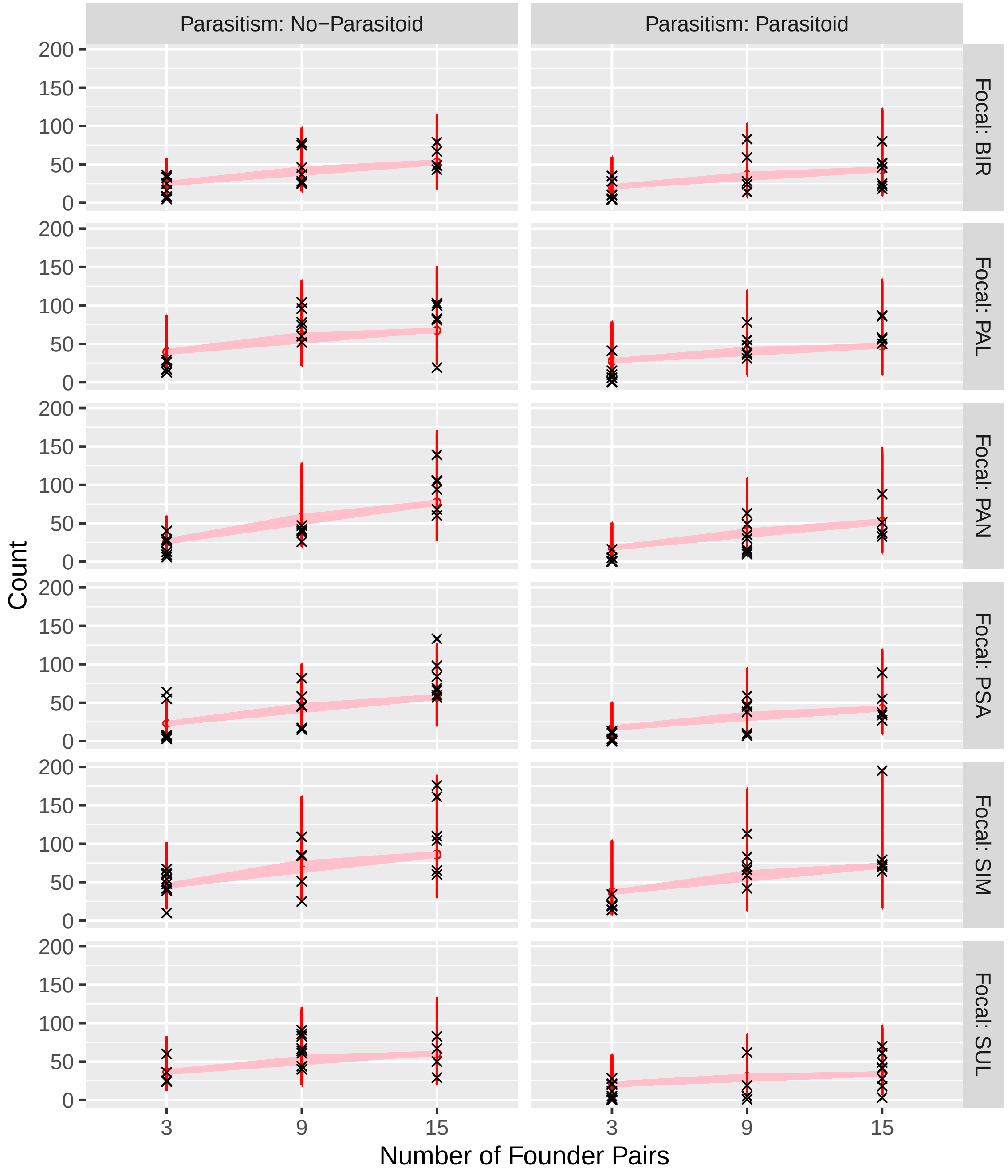


**Figure S4.8** Posterior predictions of intraspecific competition. Red dots show median predictions, red lines span central 90% of posterior distribution. Black crosses show observations.

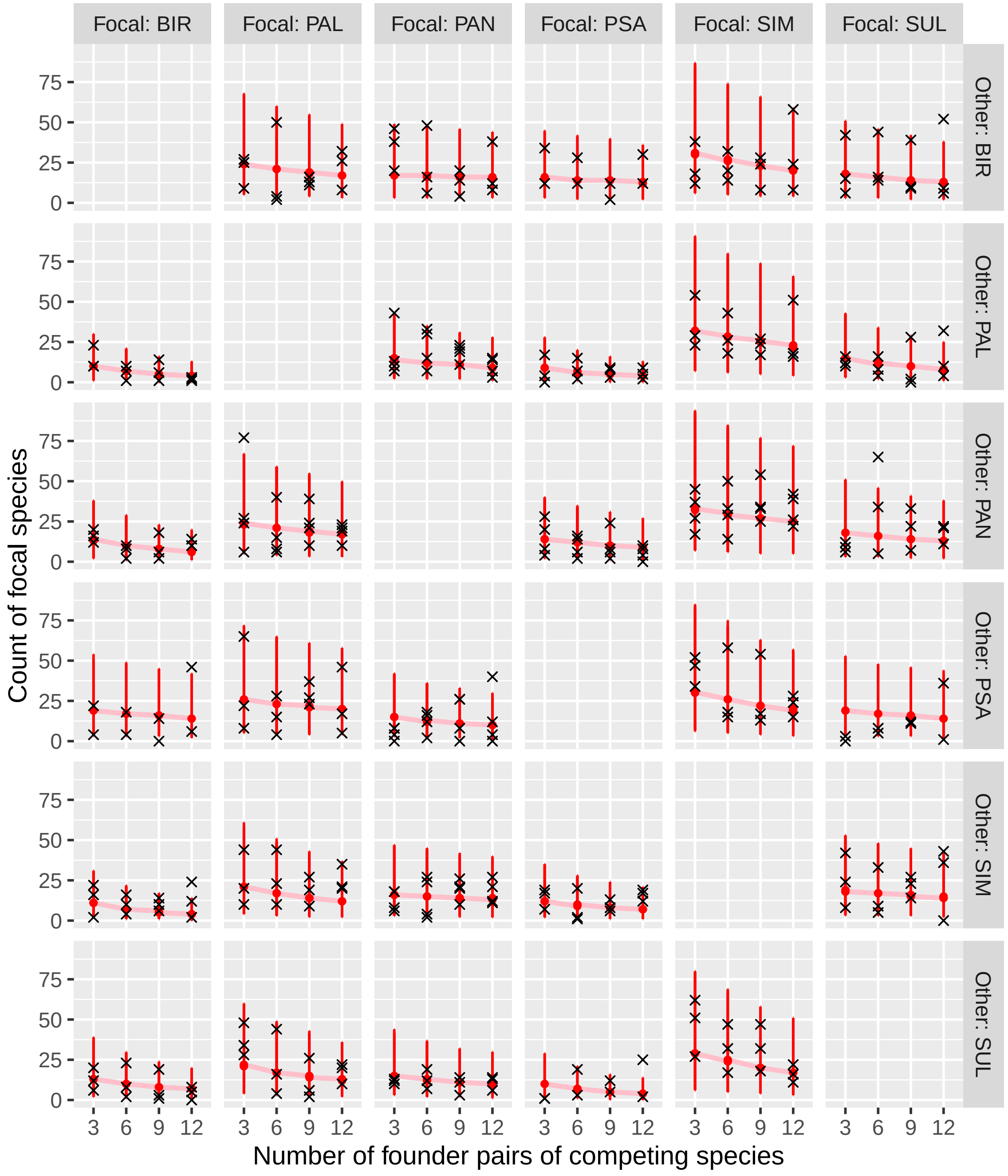


**Figure S4.9** Posterior predictions of interspecific competition in the presence of the parasitoid. Red dots show median predictions, red lines span central 90% of posterior distribution. Black crosses show observations.

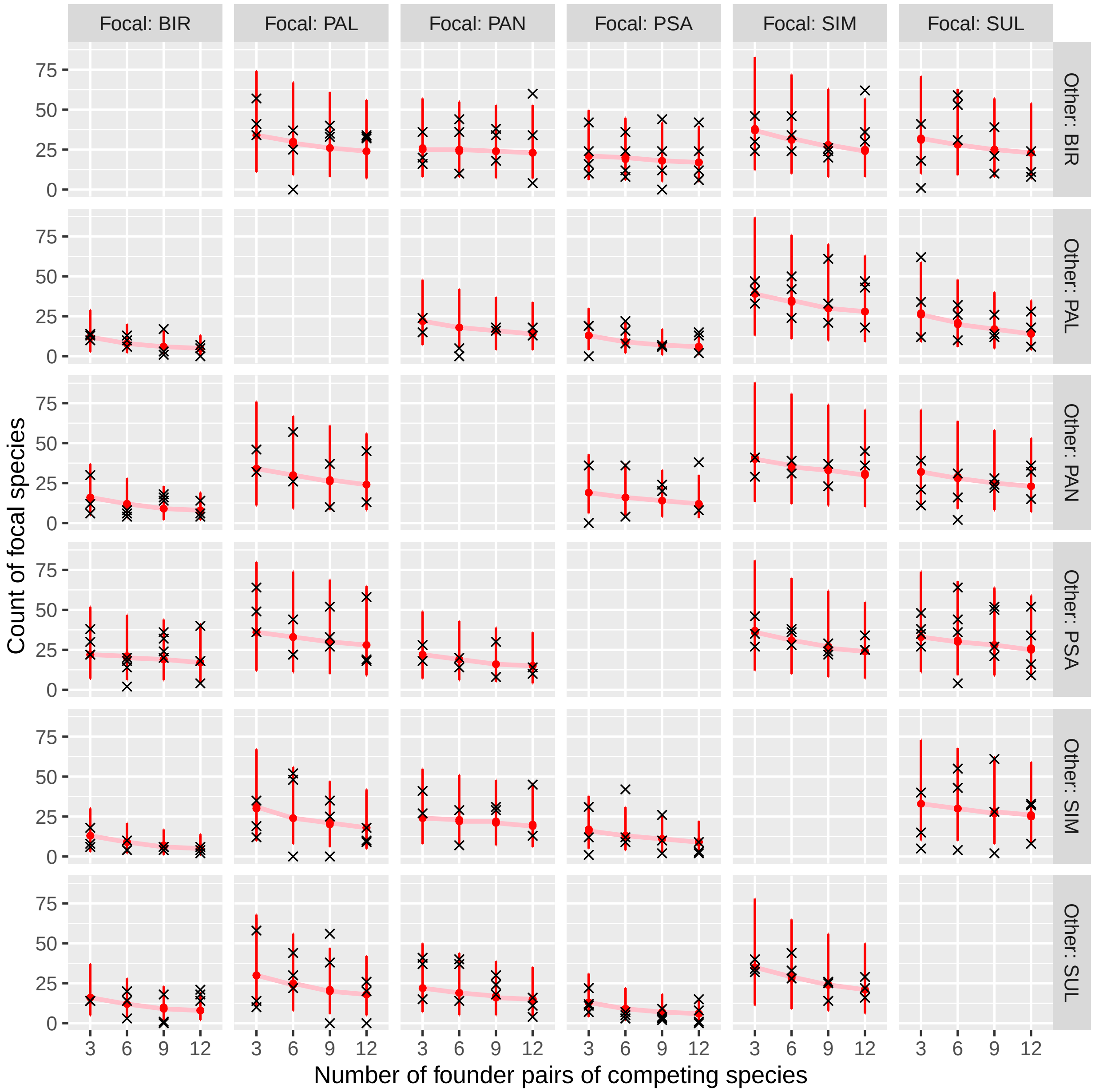


**FigS4.10** Posterior predictions of interspecific competition without the parasitoid. Red dots show median predictions, red lines span central 90% of posterior distribution. Black crosses show observations.

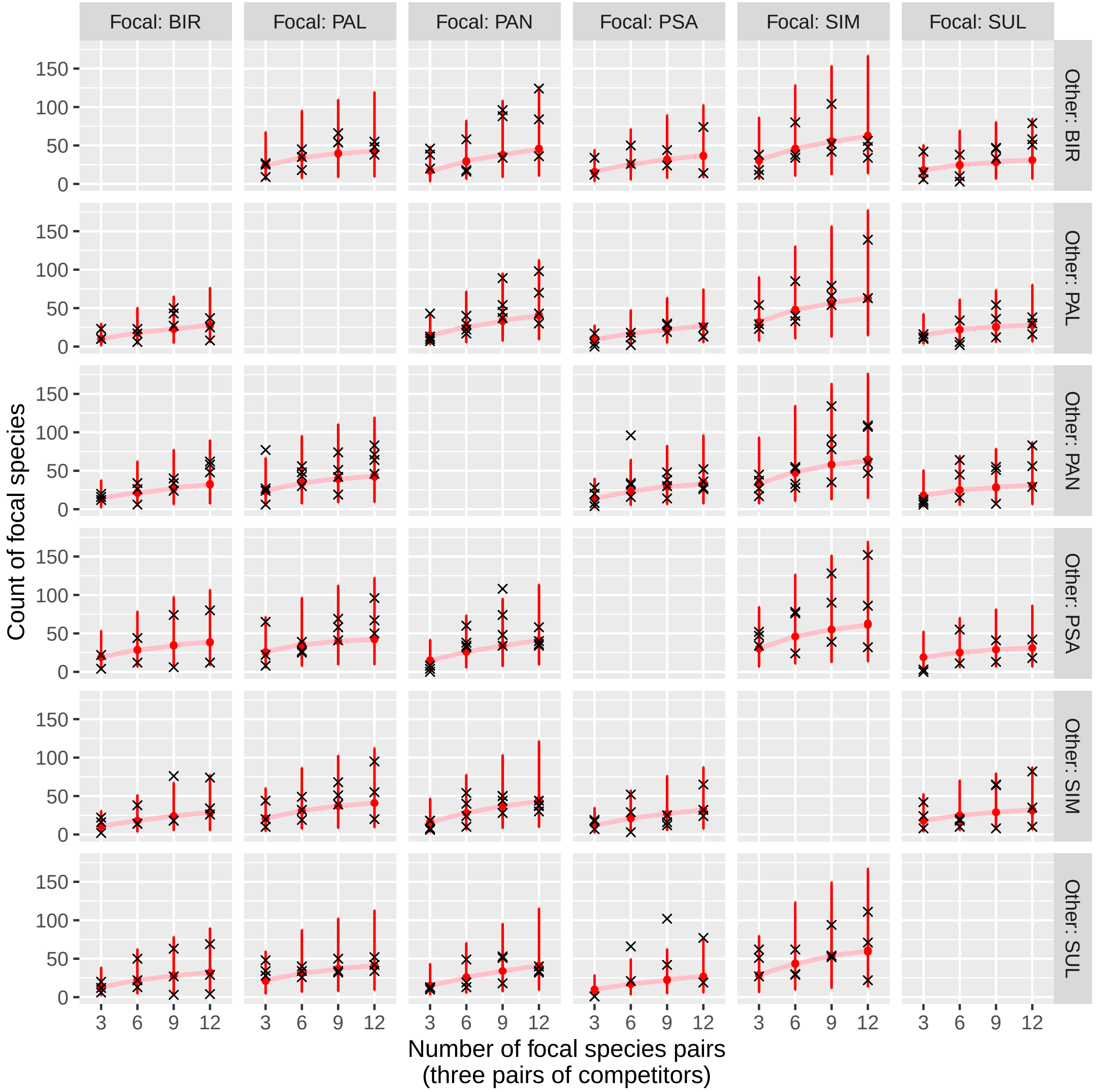


**FigS4.11** Posterior predictions covering remaining observations from the parasitoid treatment not easily displayed in previous plots, where the density of the focal species varies and the competing species is represented by 3 founder pairs. Red dots show median predictions, red lines span central 90% of posterior distribution. Black crosses show observations.

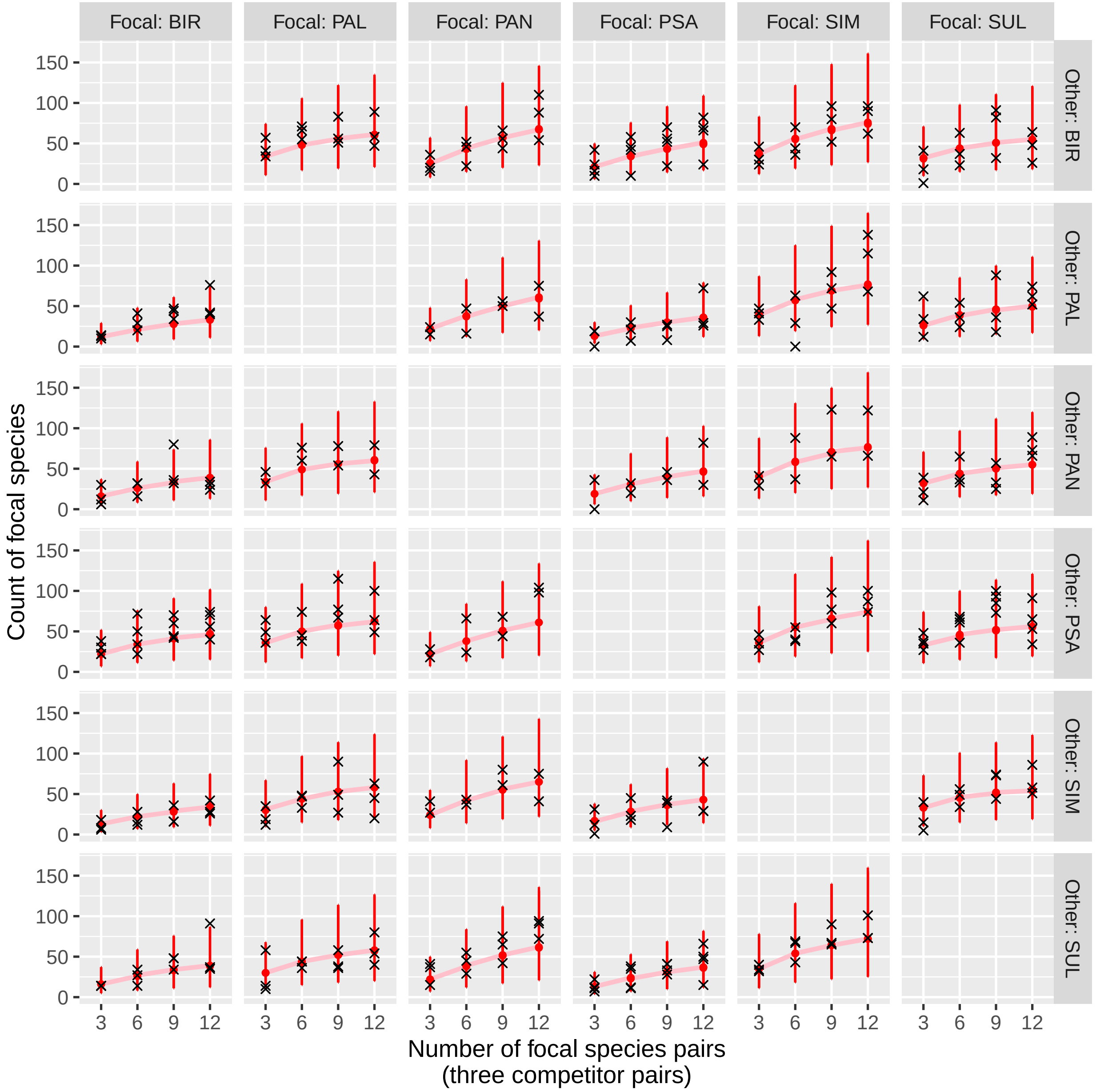


**FigS4.12** Posterior predictions covering remaining observations from the no-parasitoid treatment not easily displayed in previous plots, where the density of the focal species varies and the competing species is represented by 3 founder pairs. Red dots show median predictions, red lines span central 90% of posterior distribution. Black crosses show observations.

#### E. Maximal model

Model comparison indicates that there is little support for fitting changes in competitive coefficients due to the parasitism treatment. For completeness, we explore this 'maximal' model here. As discussed in the main text, it is possible that an intermediate model, where some competition terms vary, but others are fixed. Fig. S4.13-14 show the divergences in fitted $\alpha$ values between treatments. On average, competition coefficients are larger under parsistism than without.

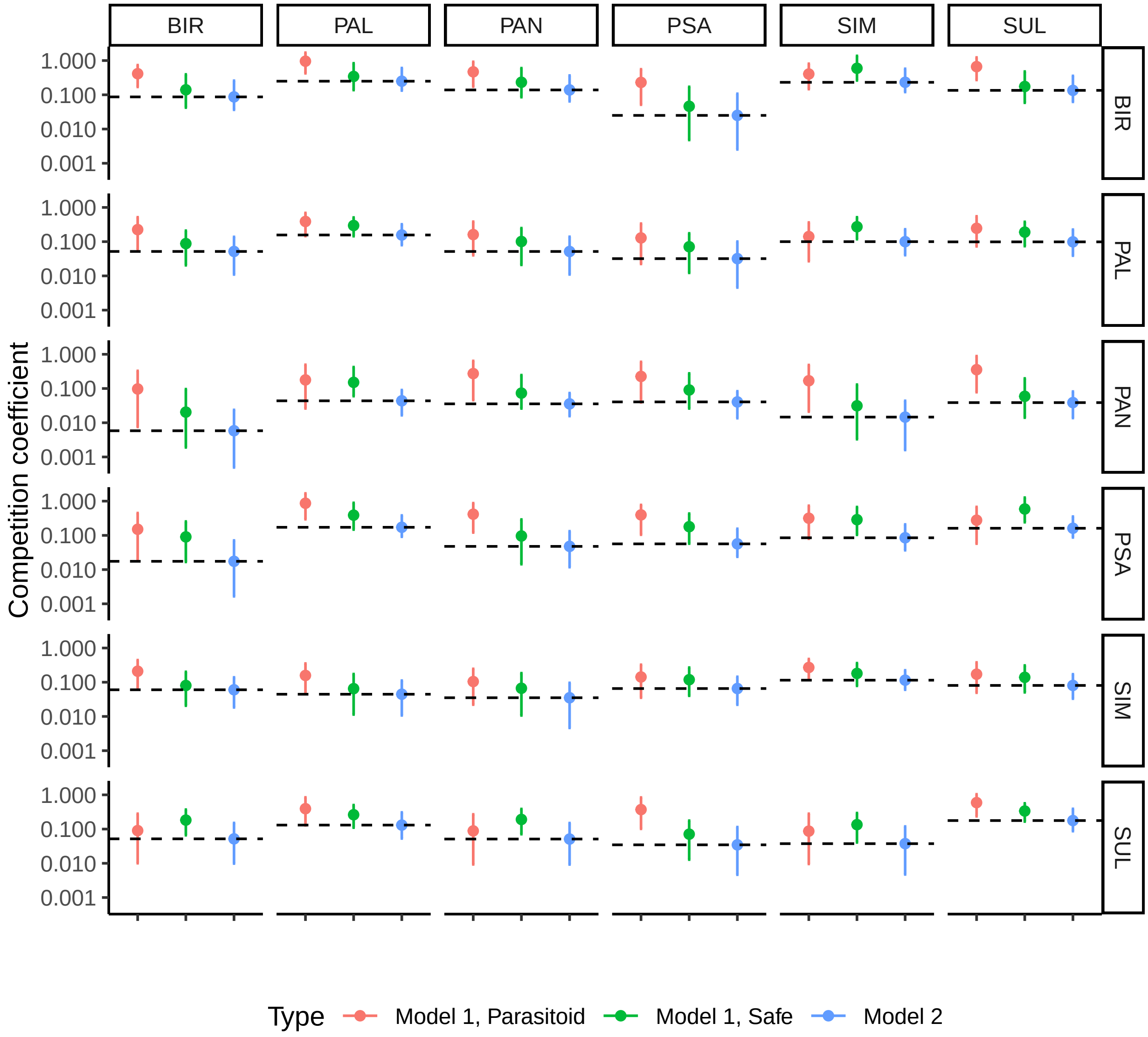


**Fig S4.13** Comparison of fitted $\alpha$ values between Model 2 (where a single $A$ matrix is fitted across both treatments) and Model 1 (where a separate $A$ matrix is fitted for each treatment). Dotted line shows median value for Model 2 to allow easier comparison. Error bars show 90% CI.

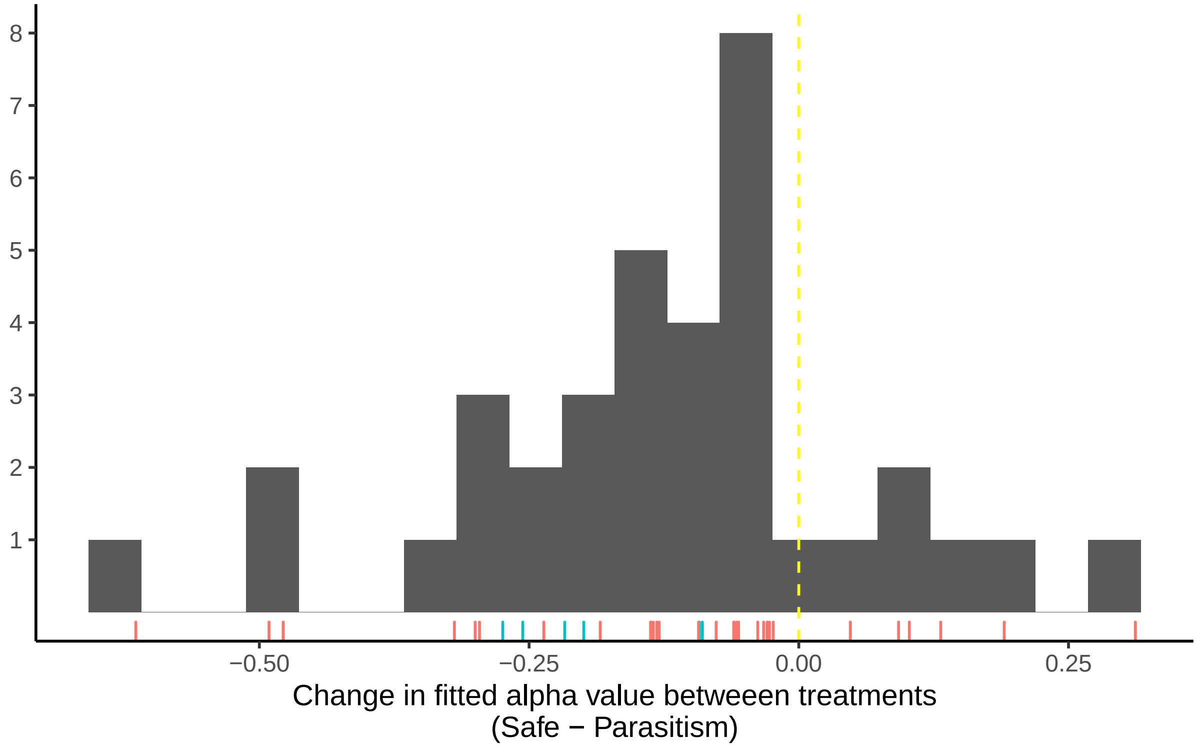


**Figure S4.14.** Distribution of shifts in median fitted $\alpha$ value between treatments in Model 1. Intraspecific terms are shown in blue, interspecific in red.')
